# Longitudinal Single-Cell RNA-seq Profiling of Lung Cell Phenotypes, Signaling, and Cross-talk During Fibrosis Resolution

**DOI:** 10.64898/2026.04.06.716772

**Authors:** Jennifer Speth, Vivian T. Wong, Steve D. Guzman, Yang Liu, Natalie M. Walker, Rachel L. Zemans, Timothy S. Blackwell, Carlos A. Aguilar, Marc Peters-Golden, Sean M. Fortier

**Author notes:** <u>Corresponding Author</u>: Sean M. Fortier, MD 6301B MSRB III, 1150 W. Medical Center Dr. Ann Arbor, MI 48109-5642.

## Abstract

Resolution of fibrosis following lung injury is distinguished from persistent/progressive parenchymal scarring through the timely clearance of aberrant cell types, removal of excess collagens, and regeneration of alveolar structure. The requisite signaling pathways, cellular cross-talk, and phenotypic shifts associated with, and required for, resolution of established lung fibrosis have not been well characterized. To address this critical knowledge gap, we performed longitudinal single-cell RNA sequencing of whole mouse lung digests obtained during spontaneously resolving fibrosis. We observed a putatively pro-fibrotic macrophage population emerge during peak fibrosis and undergo partial clearance during resolution. Our study also revealed conspicuous shifts in well-established pathways associated with tissue repair and fibrosis among immune, mesenchymal, and epithelial cells during spontaneous resolution. In addition to a decline in pro-fibrotic driver pathways, the putative anti-fibrotic pathways cAMP, HGF/MET, and TWEAK were enriched in several cell types during spontaneous resolution. CellChat analysis was used to predict the cellular senders and recipients of each pathway and characterize their longitudinal changes. Our characterization of the cellular and molecular dynamics in whole lungs during spontaneous fibrosis resolution provides a foundation for the identification of endogenous pathways that might be leveraged to treat pulmonary fibrosis.

## INTRODUCTION

Pulmonary fibrosis (PF) is a pathologic condition in which the lung’s normal wound-healing response goes awry. Because available antifibrotic drugs fail to halt or reverse lung scarring, this disease continues to exert enormous morbidity and mortality. For decades, PF research has primarily focused on characterizing the cellular and molecular basis for lung fibrogenesis, with much less attention to the cellular signaling pathways and phenotypic shifts operative in normal repair. As individuals with PF often exhibit substantial parenchymal scarring at the time of diagnosis (1), identification of these endogenous processes and cellular events associated with fibrosis resolution following injury in healthy lungs has the potential to provide novel targets to treat and possibly reverse PF.

Though the specific mechanisms that orchestrate fibrosis resolution are not well understood, it ultimately involves the removal of excess collagen/matrix proteins and a return of cellular and molecular homeostasis enabling restoration of a normal gas exchange surface (2). This, in turn, depends upon phenotypic/functional alteration or clearance of aberrant specific cell types, namely fibroblasts, epithelial cells, and macrophages. A well-known experimental example of spontaneous fibrosis resolution following lung injury is the single-dose bleomycin (bleo) model. Following intrapulmonary bleo administration, young healthy mice reliably develop fibrosis by day 14 (peaking by day 21) and exhibit near complete resolution by day 63 (3–6). Although cell-specific genetic perturbations have been shown to abolish spontaneous resolution (4, 6–9), the cross-talk between lung cells, their subtypes, and the operative pathways during resolution remain poorly characterized.

Single-cell RNA sequencing (scRNA-seq) has provided new and rich insights into the cellular heterogeneity among lung cell types in a variety of disease models. Such studies of fibrotic human and mouse lungs have revealed mesenchymal, epithelial, and immune cell subtypes that play key roles in the development of fibrosis (10–19). By contrast, we are not aware of any comprehensive single-cell investigations that have simultaneously profiled all lung cells *in situ* in an unselected manner in order to evaluate dynamic shifts in cellular phenotype, key regulatory pathways, and intercellular cross-talk throughout the process of spontaneous resolution. A holistic single-cell approach without prior cell enrichment has the potential to provide an unbiased assessment to uncover key insights into tissue remodeling, identify pathways associated with fibrosis resolution, and reveal potential targets for intervention to promote lung regeneration.

We performed longitudinal scRNA-seq on whole lung digests from young healthy mice to characterize the shifts in cellular composition, phenotype, and crosstalk during spontaneous fibrosis resolution. Specifically, mice were treated with a single dose of intrapulmonary bleo (day 0) and harvested for whole lung digest scRNA-seq on days 21, 42, and 63 – time points that we have previously established as representing peak fibrosis, ∼50% (mid) resolution, and ∼95% (full) resolution, respectively (6, 7). Our study provides the first descriptive single-cell transcriptomic analysis of whole lung tissue during spontaneous fibrosis resolution.

## METHODS

### Animal Studies

Female C57BL/6 mice homozygous for a Cre-inducible Rosa26 tdTomato lineage tag were bred with male mice heterozygous for *Cre* coupled to a tamoxifen inducible (ERT2) *Col1a2* promoter. The *Col1a2^CreERT2+/0^*;*R26^tdT+/0^* progeny were aged to 12 weeks and treated with o.p. saline or bleo (1.0 U/kg, Millipore Sigma, B5507) at day 0 as described previously (9). Mice were sacrificed on days 21, 42, or 63. Lungs were perfused with cold PBS, and the entire lung was harvested to generate RNA libraries for sequencing. Specifically, perfused lungs were minced followed by end over end rotation in a solution of 0.1% collagenase A (Roche; cat# 10103578001) in serum free DMEM for 30 min at 37C. Tissue pieces were further dissociated using an 18G needle, followed by straining through a 40 μm cell strainer to remove debris and undigested tissue. Cells were then pelleted by spinning at 500xg for 10 min and resuspended in PBS followed by pooling of equal numbers of male and female derived cells for scRNA-seq library preparation. In a parallel experiment, the right lower lobe was frozen in liquid nitrogen, embedded in paraffin, and subsequently sectioned to perform Masson’s trichrome staining, followed by imaging with bright-field, and whole-slide imaging with a Vectra Polaris Brightfield Scanner.

### scRNA-seq data processing and analysis

Raw sequencing data were processed using CellRanger (version 7.2.0, 10x Genomics). Sequencing reads were aligned to the *Mus musculus* reference transcriptome (mm10) using the STAR aligner. The CellRanger count pipeline was run with default parameters to generate gene expression count matrices. Raw and filtered feature barcode matrices were imported into R (version 4.3.1) using the Read10X function.

All downstream analysis was performed in R using Seurat (version 5.0.0). Four datasets, corresponding to distinct experimental time points (saline control, day 21, day 42, day 63) were preprocessed independently. Each dataset consisted of high-quality singlet cells that underwent quality control as described below.

Ambient RNA contamination was corrected using SoupX (version 1.6.2). A SoupChannel object was created from the raw and filtered matrices, and the contamination fraction was estimated from the expression profiles of empty droplets. This estimated background was then used to adjust expression counts for individual cells, and corrected count matrices were saved for downstream analysis.

Standard quality control thresholds were applied to remove low-quality cells and potential multiplets (Supplemental Figure 1B). Cells with fewer than 200 detected genes or more than 10,000 genes, fewer than 500 or more than 20,000 total RNA counts, or greater than 10% of UMIs mapping to mitochondrial genes were excluded. Filtered data were normalized using log-normalization, regressing out total UMI counts and the percentage of mitochondrial transcripts per cell. Variable features were identified using the VST method, and principal component analysis (PCA) dimensionality reduction was performed. Data were further normalized using SCTransform (glmGamPoi method) to stabilize variance across sequencing depth. The first 30 principal components were used for uniform manifold approximation and projection (UMAP) embedding and unsupervised clustering based on a shared nearest neighbor (SNN) with a resolution of 0.5.

Doublet detection was performed using DoubletFinder (version 2.0.4). The expected doublet rate was set to 7.5% based on 10x Genomics guidelines for the estimated cell recovery rate (20), and default parameters were used otherwise. Cells classified as singlets were then extracted for downstream analysis.

Cell type annotation was conducted with Clustifyr (version 1.8.0). Cluster-level expression profiles were correlated to two mouse lung single-cell reference datasets from CZ CellxGene and ScTypeDB (21, 22). Gene identifiers were harmonized to *Mus musculus* gene symbols using biomaRt. Cluster-level correlations were computed using clustify and cell type identities were assigned to each cluster. Manual review and refinement of annotations was performed as needed. Differential gene expression analysis was conducted across annotated cell types to identify key marker genes. Subsets of selected cell populations (fibroblasts, epithelial cells, and macrophages) were reanalyzed using the same Seurat workflow, with clustering resolution increased to 0.7 to capture finer substructure within lineages.

Investigation of cell-cell communication networks was performed using CellChat (version 1.6.1) on the datasets. Analyses were performed both on the full dataset and on a subset of selected cell types using metadata from the Seurat cluster identities from the SCT assay. The CellChat mouse ligand-receptor database (including non-protein interactions) was used to predict signaling interactions, with cell populations with fewer than 10 cells excluded from the analysis. Overexpressed genes and ligand-receptor interactions were identified, followed by computation of communication probabilities between cell types and pathway and centrality analyses. All CellChat analyses were run with default parameters for functions and visualization.

### Immunofluorescence Microscopy

Mouse lungs were fixed with 10% formalin, embedded in paraffin, sectioned, and mounted on glass slides. Subsequent deparaffinization, citrate antigen retrieval, blocking and staining was performed as previously described (7). Staining of fixed fibroblasts and murine lung tissue was performed with the following primary antibodies: rabbit anti-CD63 (1:500; HPA010088; Sigma) and rabbit anti-Arg-1 (1:500; 16001-1-AP; Proteintech) overnight at 4°C. They were then incubated with anti-rabbit secondary antibodies (FITC (1:200; Jackson 711-545-152) and Cy5 (1:200; Jackson 107-605-142). Mounting medium containing DAPI (Invitrogen) was used to stain nuclei. Images were obtained using a Nikon Eclipse Ti2 Inverted Confocal or an Olympus BX53 microscope.

## RESULTS

### scRNA-seq of the mouse lung during spontaneous fibrosis resolution

Previous scRNA-seq studies during spontaneous PF resolution have employed initial sorting of various prespecified lung cell populations (19, 23), To examine these dynamic changes at an unbiased whole lung cell level, we conducted scRNA-seq on unsorted lung digests harvested from mice following a single dose of intrapulmonary bleo (1.0 U/kg) at day 21 (peak fibrosis), day 42 (partial [∼50%] resolution), and day 63 (full [∼95%] resolution Figure 1A). Histopathologic analysis using Masson’s trichrome staining confirmed increased collagen deposition and fibrotic changes at day 21 compared to saline control, with histopathological evidence of fibrosis decreasing progressively through days 42 and 63 (Figure 1B) as has been previously reported (3–7, 24). We generated 4 high-quality scRNA-Seq libraries encompassing on average 2,388 genes and 7,685 unique molecular identifiers per cell (Supplemental Figure 1). After cell digestion and sequencing using the Illumina 10X and standard Seurat workflow (Figure 1C), we employed filtering to reduce the dimensionality of the datasets using Uniform Manifold Approximation and Projection (UMAP) and Louvain clustering which displayed 20 cell types (Figure 1D). Cell types were annotated using established identification markers from two mouse lung single-cell datasets (21, 22), revealing the presence of dendritic cells, T cells, B cells, NK cells, neutrophils, monocytes, interstitial macrophages (IM), alveolar macrophages (AM), vascular endothelial cells, lymphatic endothelial cells, aerocytes (aCap endothelial), general capillary (gCap endothelial) cells, alveolar type 2 (AT2) and type 1 (AT1) epithelial cells, club cells, ciliated cells, pericytes, smooth muscle (SM) cells, mesothelial cells, and fibroblasts (Figure 1D-E). At peak fibrosis, we observed modest shifts in certain immune cell populations (T cells, B cells, neutrophils, and IMs) compared to baseline (control); these changes returned to baseline levels during resolution (days 42 and 63). Notably, at whole lung resolution, no substantial changes were detected in the proportions of mesenchymal (fibroblasts, pericytes, and SM cells) or epithelial (AT1, AT2, club, and ciliated cells) populations throughout disease progression and resolution (Figure 1F).

**Figure 1:**
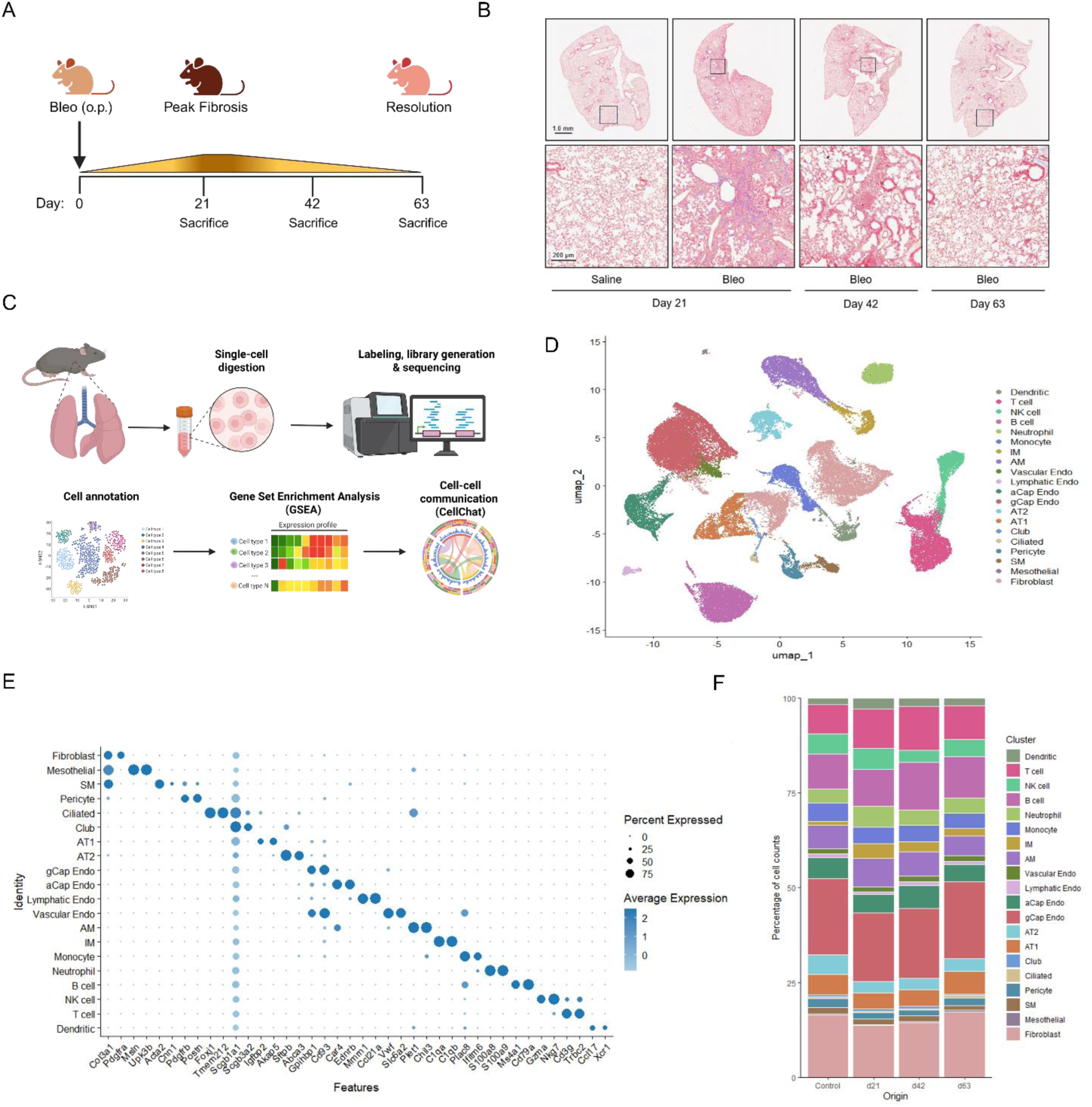
Longitudinal scRNA sequencing of the mouse lung during spontaneous fibrosis resolution. (A) Schematic illustrating the single-dose bleomycin protocol. (B) Masson’s trichrome staining of right lower lobe sections demonstrating spontaneous fibrosis resolution. (C) Schematic detailing generation of whole lung digest, library prep, single-cell RNA sequencing and analysis. (D) Compiled Uniform Manifold Approximation and Projection (UMAP) plot with cell annotation of whole lung digests at each time point condition. (E) Dot plot displaying top gene expression for each annotated cell type in D. (F) Cluster graph displaying longitudinal cell type composition.

### Lung immune, mesenchymal, and epithelial cells exhibit bidirectional signaling and pathway enrichment during fibrosis resolution

To investigate cell communication dynamics, we employed CellChat analysis (25) to quantify both the relative incoming and outgoing signaling strengths for each cell population, as well as the number of interactions between different cell types.

AMs, consistent with their central role in orchestrating lung immune responses, demonstrated high interaction strength for both incoming and outgoing signals (Figure 2A). In contrast, other innate immune cells (dendritic cells, monocytes, and neutrophils) and adaptive immune cells (T and B cells) displayed substantially lower levels of outgoing signaling. Fibroblasts, AT1 and AT2 epithelial cells, and IMs exhibited moderate levels of both incoming and outgoing signals. We found that both IM and AM populations primarily communicate with each other as well as with mesenchymal cell populations (Figure 2B). This pattern is also mirrored in fibroblasts, which demonstrate heightened signaling with IMs and AMs, as well as other mesenchymal cell populations.

**Figure 2:**
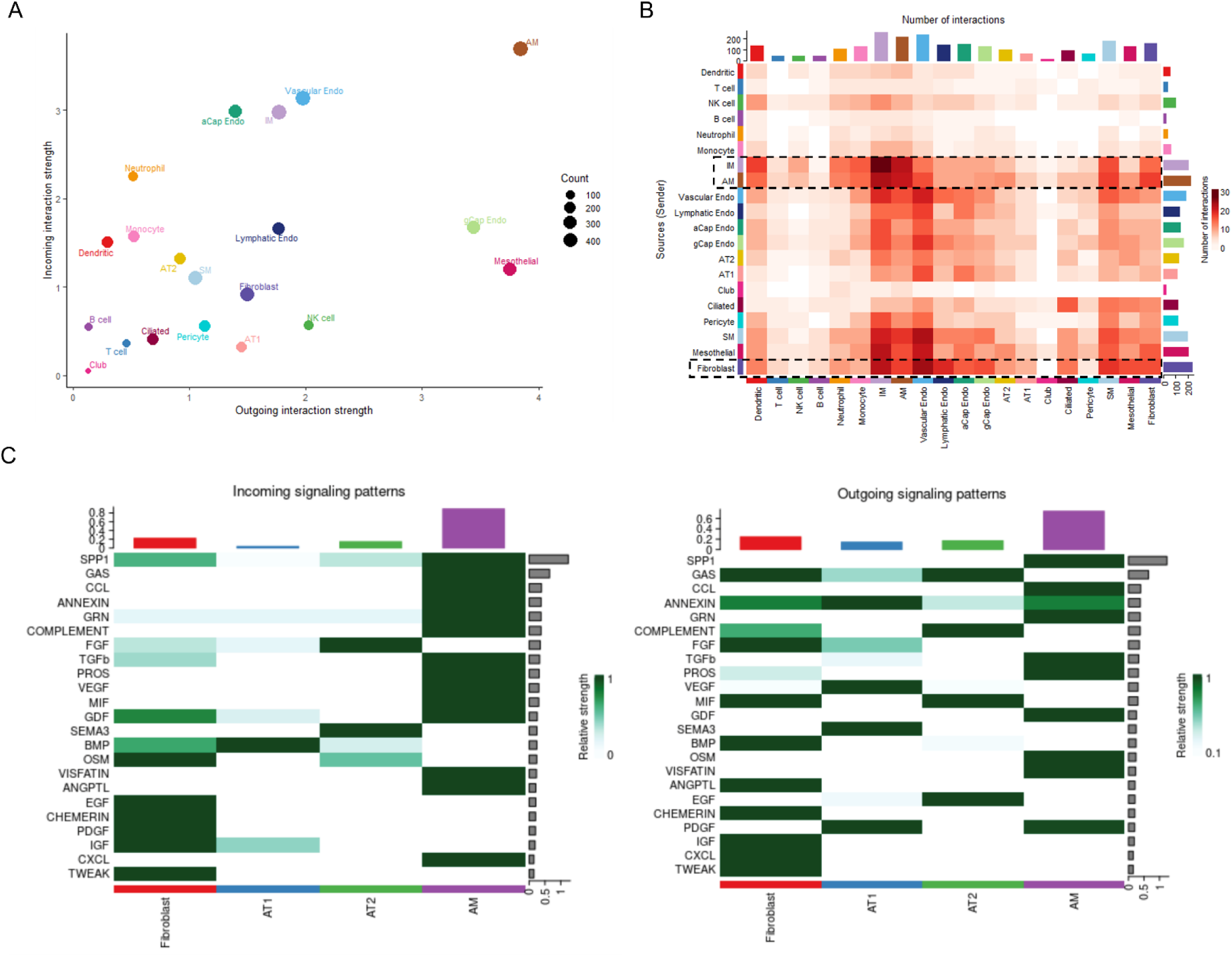
Relative intercellular signaling strength, cross-talk, and pathway enrichment among lung cells during fibrosis resolution. (A) Dot plot depicting relative incoming and outgoing interaction strength by all annotated cell types. (B) Heat map depicting relative number of interactions between sender cell types (y-axis) and receiver cell types (x-axis). (C) Heat map of top enriched pathways among fibroblast, type I (AT1), type II (AT2) alveolar epithelial cells, and alveolar macrophages (AMs).

Focusing on cell types typically considered most relevant to disease pathology and their specific communication patterns, we conducted a detailed analysis of incoming and outgoing signaling pathways among AMs, AT1 cells, AT2 cells, and fibroblasts (Figure 2C). Consistent with their role as facilitators of fibrosis, fibroblasts received incoming signals that activate pathways known to be pro-fibrotic including osteopontin (SPP1) (26), fibroblast growth factor (FGF) (27), platelet-derived growth factor (PDGF) (28, 29), and transforming growth factor-beta (TGFβ) (27). They also produced outgoing pro-inflammatory signals such as migration inhibitory factor (MIF), complement, C-X-C motif chemokines (CXCLs), and oncostatin M (OSM), supporting the idea that fibroblasts can adopt an inflammatory phenotype that may contribute to the fibrotic process (30, 31). AMs also demonstrated increased fibrotic signaling, characterized by the marked strength of their outgoing SPP1, TGFβ, and PDGF signaling as well as inflammatory C-C motif chemokines (CCL) signaling. AT1 alveolar epithelial cells produced various growth factors including FGF, VEGF, and PDGF. AT2 cells were found to be a source of complement and an additional source of MIF. AT2 cells have been shown to promote lung fibrosis through MIF-dependent cross-talk with lung macrophages (32). Notably, fibroblasts, AT2, and to a lesser degree AT1 cells, were all found to be a source of Growth Arrest–Specific (GAS) signaling with AMs as a likely recipient cell type. Among this family of ligands, GAS6 has been reported to exert anti-inflammatory effects in lung macrophages (33). Interestingly, pathways associated with both “pro-resolution” and “pro-fibrotic” activity (depending on cell type, context, and the organ involved) were also enhanced; these include TNF-related weak inducer of apoptosis (TWEAK) (34) (fibroblasts), growth differentiation factor (GDF) (35, 36) (fibroblasts and AT1), and bone morphogenic protein (BMP) (37) (fibroblasts and AT1) (Figure 2D).

Though not specifically identified previously in the context of fibrosis resolution, these findings are consistent with many well-characterized pro- and anti-fibrotic and -inflammatory pathways operative within immune, mesenchymal, and epithelial cells during lung fibrogenesis and its spontaneous resolution. Several instances of paired incoming and outgoing signaling among different cell types suggest the potential for cross-talk among them. To gain a deeper understanding of how enrichment of these and other pathways might orchestrate resolution, we sought to determine which macrophage, fibroblast, and epithelial cell subtypes send and receive the relevant corresponding signals and to determine their patterns of expression throughout resolution. Broadly assessing overall signaling strength within cell types longitudinally, we found that the magnitude of incoming and outgoing signaling was reduced during peak fibrosis compared to saline-treated control lungs, with further relative decline in signaling strength by day 63 (Supplemental Figure 2). We next sought to subcluster macrophages, mesenchymal cells, and epithelial cells and analyze them longitudinally.

### Fibrosis-associated macrophages alter their signaling pathways and are largely cleared during spontaneous fibrosis resolution

To more precisely characterize changes in lung macrophage subsets during fibrosis and resolution, clusters identified as “AMs” and “IMs” from the global UMAP analysis in Figure 1 were further subclustered. This approach revealed four distinct populations based on established marker genes (Figure 3A-B). Tissue-resident AMs (TRAMs) exhibited high expression of *SiglecF*, *Ear1*, *Car4*, *Cd11c* (*Itgax)* and *Fabp5*. Two IM populations were distinguished from TRAMs by elevated levels of *CD11b (Itgam)*, *CX3CR1*, and *C1qa*, with one distinct population expressing the vascular-associated marker *Lyve1*. Finally, a cluster co-expressing both IM and TRAM markers appeared at peak fibrosis consistent with those described in previous studies of bleo-induced lung injury (10, 38, 39). Furthermore, this population expressed many of the hallmark genes associated with a pro-fibrotic lung macrophage phenotype including *Arg1* (39), *Spp1* (40, 41), *Trem2* (40, 42), *CD63* and *Fabp5* (40) (Figure 3D, Supplemental Figure 3B). We thus labeled this cluster fibrosis-associated macrophages (FAMs) due to their appearance at peak fibrosis (day 21) and likewise enrichment in pro-fibrotic genes. Notably, cell proportion analysis indicated a reduction in FAMs during the resolution phase (Figure 3C, Supplemental Figure 3A). To confirm their presence *in vivo*, we performed immunofluorescence (IF) microscopy on lung sections from saline-treated mice and those harvested on day 21 after bleo treatment. IF staining revealed the presence of Arg1+CD63+ macrophages at day 21, consistent with their detection by scRNA-seq (Figure 3D-F).

**Figure 3:**
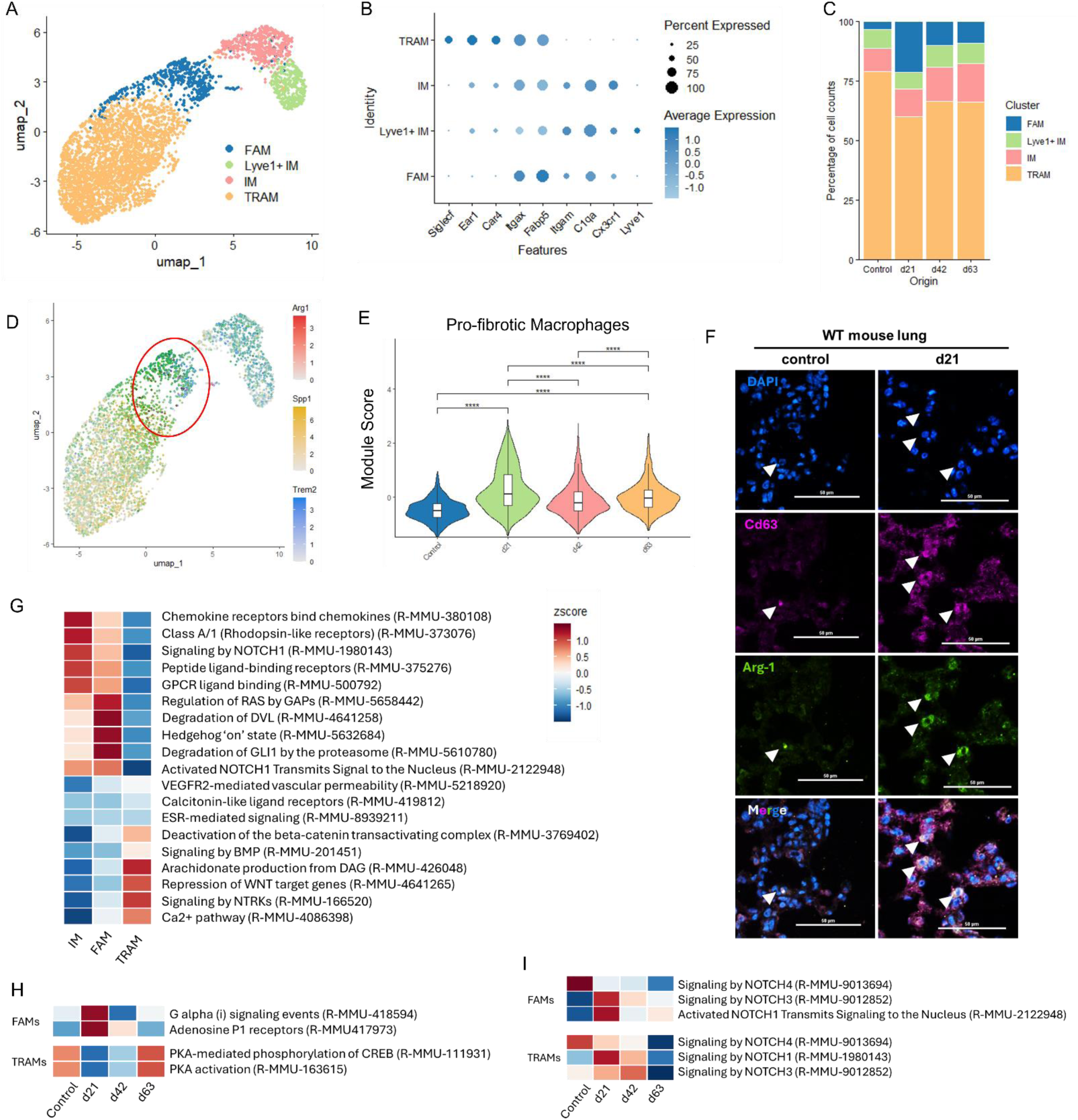
Fibrosis-associated macrophages alter their signaling pathways and are largely cleared during spontaneous fibrosis resolution. (A) UMAP of macrophage subclusters displaying tissue resident alveolar macrophages (TRAMs), interstitial macrophages (IMs), Lyve1+ IMs, and transitional macrophages (FAMs). (B) Dot plot of representative markers for each macrophage subtype. (C) Cluster graph displaying longitudinal cell type composition of macrophage subtypes. Feature plot displaying pro-fibrotic FAM-signature genes (Arg1, Spp1, Trem2) (D) and violin plot of module scores assigned to Arg1+ Spp1+ Trem2+ FAM subtypes (E) over time. (F) Representative immunofluorescent microscopy images of CD63 and Arg1 staining in FAMs from control and d21 fibrotic lungs. (G) Gene Set Enrichment Analysis (GSEA) of top enriched pathways within macrophage subclusters. H & I: Longitudinal GSEA of selected pathways related to cAMP (H) and NOTCH signaling (I) and related pathways within FAM and TRAM subsets.

Gene set enrichment analysis (GSEA) of the highest enriched pathways within each macrophage subtype demonstrated that FAMs were highly enriched for pathways related to sonic hedgehog (SHH), Notch1, and non-canonical Wnt signaling (as indicated by DVL degradation), further supporting their pro-fibrotic/pro-inflammatory phenotype. Interestingly, FAMs also demonstrated similar enhancement in the pathways enriched in IMs including those related to GPCR ligand binding – receptors canonically regulating calcium or cAMP signaling. TRAMs were enriched for arachidonate/eicosanoid and anti-Wnt signaling pathways (Figure 3G).

We next analyzed the enrichment of these pathways within macrophage subtypes over time to determine when, and to what relative degree, they are enriched during peak fibrosis and resolution. A parent reactome GSEA analysis of GPCR signaling within macrophage subtypes revealed that FAMs are highly enriched in Gαi and adenosine P1 activation (Supplemental Figure 3C). Gαi functions to inhibit adenylyl cyclase and thus reduce intracellular cAMP generation. Adenosine P1 receptors are coupled to G-proteins that can either inhibit or activate adenylyl cyclase. Longitudinally, adenosine P1 receptors, and Gαi enrichment generally, was found to be enriched in FAMs at day 21 (peak fibrosis) yet declined during resolution (Figure 3H).

Further analysis revealed enrichment of the inhibitory A3 receptor in FAMs at day 21 along with stimulatory A2 receptors at day 21 with decline throughout resolution (Supplemental Figure 3E). Complementary to these findings in FAMs, TRAMs exhibited a high degree of PKA activation (a downstream target of cAMP) and PKA-mediated phosphorylation of CREB in control lungs, which diminished at peak fibrosis and then recovered during resolution (Figure 3H).

Both TRAMs and FAMs demonstrated enrichment for the inflammation-associated NOTCH1 and fibrosis cross-talk associated NOTCH3 signaling pathways at day 21 with substantial decline in each by day 63 (Figure 3I). Wnt5a was found to be the only significant ligand enriched within FAMs and TRAMs among this broad signaling pathway. In FAMs, Wnt5a signaling was highly enriched at day 21 while only modestly enriched in TRAMs at this time point compared to baseline (Supplemental Figure 3D).

### Fibroblasts exhibit dynamic pro- and anti-fibrotic phenotypes and signaling during spontaneous fibrosis resolution

Given the central role of mesenchymal cells in the pathogenesis of fibrosis – primarily through their promotion of excessive extracellular matrix deposition – we sought to more precisely delineate the specific contributions of distinct fibroblast subtypes to the resolution of fibrosis. Clusters annotated as “fibroblast,” “pericyte,” “SM”, and “mesothelial cells” from the global UMAP analysis were further subclustered and subjected to reanalysis. Within the fibroblast population, we identified peribronchial, adventitial, and alveolar fibroblast subtypes, distinguished by the expression of *Hhip*, *Pi16*, and *Scube2*, respectively (43) (Figure 4A-B). As previously noted, the proportions of these fibroblast subtypes amongst the overall mesenchymal compartment remained largely unchanged throughout the fibrosis resolution process in our whole lung digests (Figure 4C). Prompted by recent studies indicating that pathologically relevant “fibrotic” fibroblast subsets may exist within the alveolar fibroblast population (19, 43), we subclustered alveolar and adventitial fibroblasts based on their elevated expression levels of the fibrotic markers *Col3a1*, *Col1a1* and *Pdgfra* (Figure 4D). The presence of a tdTomato lineage tag in mice subjected to sequencing helped to further confirm the importance of the alveolar and adventitial populations given its coupled expression to a *Col1a2* promoter (Figure 4D, Supplemental Figure 4A).

**Figure 4:**
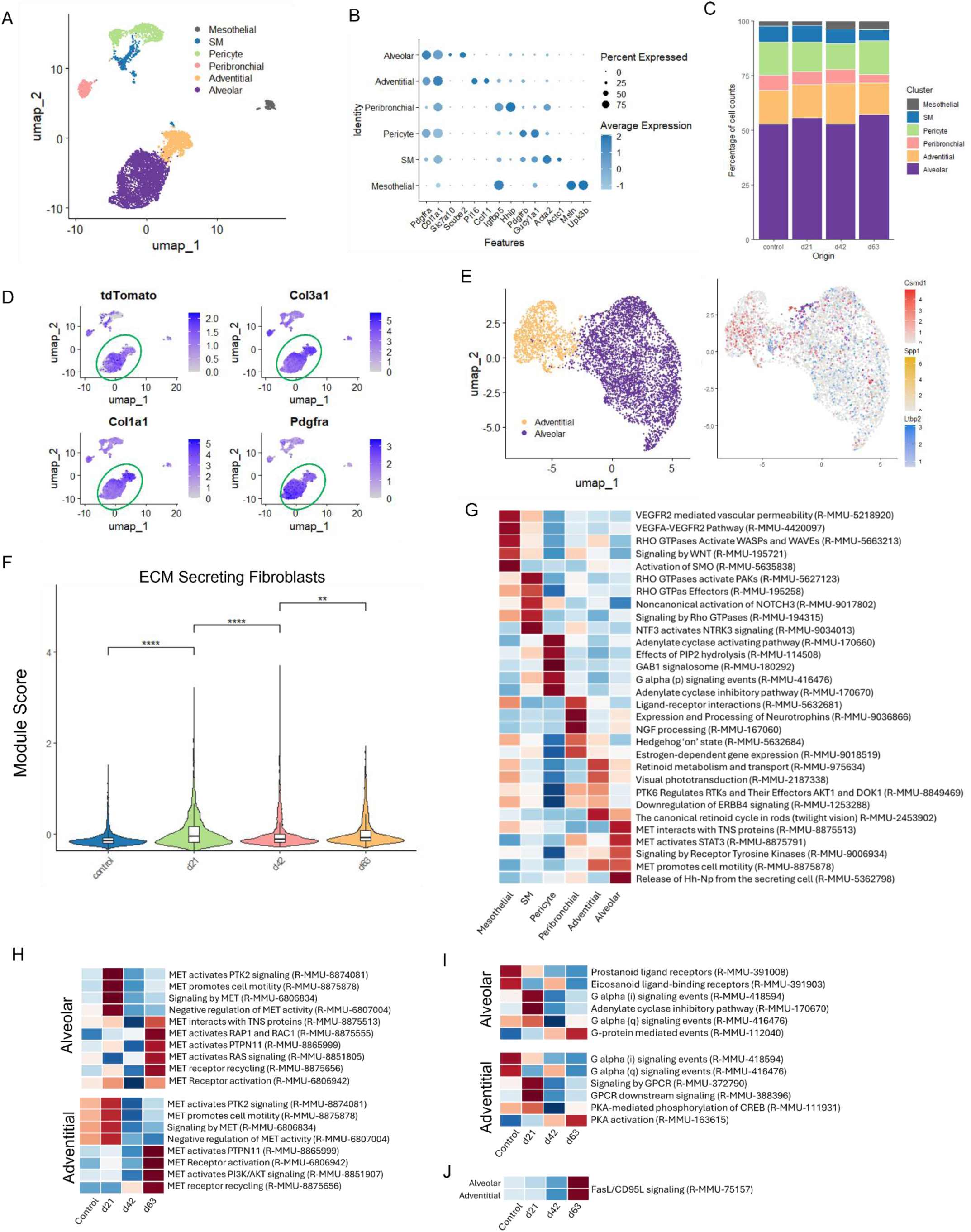
Fibroblasts exhibit dynamic pro- and anti-fibrotic phenotypes and signaling during spontaneous fibrosis resolution. (A) UMAP of mesenchymal cell subclusters. (B) Dot plot of representative markers for each mesenchymal subset. (C) Cluster graph displaying longitudinal cell type composition of mesenchymal subtypes. (D) Feature plots demonstrating relative expression of tdTomato lineage tag and fibroblast associated genes among mesenchymal cell subsets. (E) Feature plot of “ECM-secreting” fibroblast subtype within alveolar and adventitial subclusters. (F) Violin plot of module scores assigned to Csmd1+ Spp1+ Ltbp2+ ECM-secreting fibroblasts over time. (G) GSEA of top enriched pathways among mesenchymal cells. H-J: Longitudinal GSEA of selected HGF/MET (H), cAMP (I), and FasL/CD95L (J) signaling and related pathways in alveolar and adventitial fibroblasts.

We identified a distinct subpopulation previously characterized as an “ECM-secreting” fibroblast phenotype (19), as evidenced by elevated expression of *Csmd1*, *Spp1*, and *Ltbp2*. This “*Csmd1*+” population was noted within the alveolar and adventitial fibroblast compartments (Figure 4E) as well as peribronchial and smooth muscle populations but absent in pericytes (Supplemental Figure 4B). Within the alveolar and adventitial compartments, the proportion of these cells increased substantially at day 21, before declining close to baseline levels by days 42 and 63 (Figure 4F). Within the peribronchial compartment, a similar pattern of enrichment and reduction to baseline control levels was observed (Supplemental Figure 4C). Surprisingly, this ECM-secreting population did not express the pro-fibrotic marker *Cthrc1* (data not shown), which has previously been proposed as a marker of pathologic fibroblasts (43).

We next performed GSEA to determine the highest enriched pathways in each fibroblast subtype. As expected, canonical pro-fibrotic pathways involved in TGFβ, NOTCH, and hedgehog signaling were enriched in alveolar and adventitial fibroblasts at peak fibrosis (Supplemental Figure 4D). Notably, we found enhancement of MET signaling – also known as hepatocyte growth factor receptor – within the alveolar fibroblast compartment. This pathway has been shown to be anti-fibrotic in the context of pulmonary fibrosis (44, 45). Similar to the macrophage population, cAMP-signaling pathways – specifically adenylyl cyclase activation or inhibition – were found to be enriched in pericytes (Figure 4G). The calcium-signaling GPCR coupled G-protein Gαq was also found to be enriched in pericytes. Further investigation into a previously described subpopulation of putative “anti-fibrotic” fibroblasts, characterized by the expression of *Cd248*, *Pi16*, and *Igfbp5* (19), revealed no significant changes in their population frequency at any timepoint (data not shown). This observation suggests that, as seen previously, more subtle alterations in cell signaling dynamics may play a more substantial role in the resolution of fibrosis than changes in cellular abundance.

Longitudinal analysis of MET and GPCR-related pathways was performed to assess their relative enrichment throughout resolution. Within alveolar and adventitial fibroblasts, MET was found to be negatively regulated and associated with PTK2 (FAK) during peak fibrosis while its activation and interaction with various downstream mediators was enriched during resolution (Figure 4H); this is consistent with its role as an anti-fibrotic signal. With regards to GPCR-mediated signaling, both cAMP and calcium-related pathways demonstrated dynamic enrichment during peak fibrosis and throughout resolution in alveolar and adventitial fibroblast populations. Prostanoid and eicosanoid receptors – that often signal via cAMP – were found to be enriched in alveolar fibroblasts from saline-treated mice with reduced enrichment at peak fibrosis and throughout resolution. Additionally, Gαi and adenylyl cyclase inhibitor signaling were enriched at peak fibrosis in both subtypes. However, PKA activation and PKA-mediated phosphorylation of CREB were enriched at day 63 in adventitial fibroblasts (Figure 4I). In both alveolar and adventitial fibroblasts, Gαq signaling was highly enriched at peak fibrosis and reduced throughout resolution. Importantly, FasL signaling was enriched by day 63 in both alveolar and adventitial fibroblasts (Figure 4J) consistent with possible apoptotic clearance of these cells.

### Lung epithelial cells demonstrate enrichment for pro-regenerative and pro-resolution pathways during spontaneous fibrosis resolution

Epithelial cells are highly susceptible to injury from environmental exposures, infections, and inflammatory processes. Injury-induced dysfunction of this cell population serves as a key driver in the development and progression of pulmonary fibrosis (46–49). While their role in the early inflammatory stages of pulmonary fibrosis has been extensively investigated, their status during the resolution phase of fibrosis is poorly understood. To address this gap in knowledge, we analyzed epithelial cell subtypes and evaluated their potential contributions to the mechanisms underlying fibrosis resolution. Clusters corresponding to “AT1,” “AT2,” “ciliated,” and “club” epithelial cell types were subclustered from the global UMAP and subsequently reanalyzed using established cell-specific markers. This approach uncovered a “transitional” population distinguished by expression of *Krt8*, *Sfn* and *Cldn4* (49) (Figure 5A-C; Supplemental Figure 5A), which was closely aligned with both AT1 and AT2 clusters. This transitional subset was only evident in bleo-treated mice and demonstrated its highest proportion at peak fibrosis (day 21), with its abundance diminishing by days 42 and 63 (Figure 5C and Supplemental Fig 5A). Aside from this shift of transitional epithelial cells, the relative proportions of epithelial cells demonstrated a decline in AT1 cells and mild increase in club cells at peak fibrosis with relative stability throughout resolution (Figure 5C).

**Figure 5:**
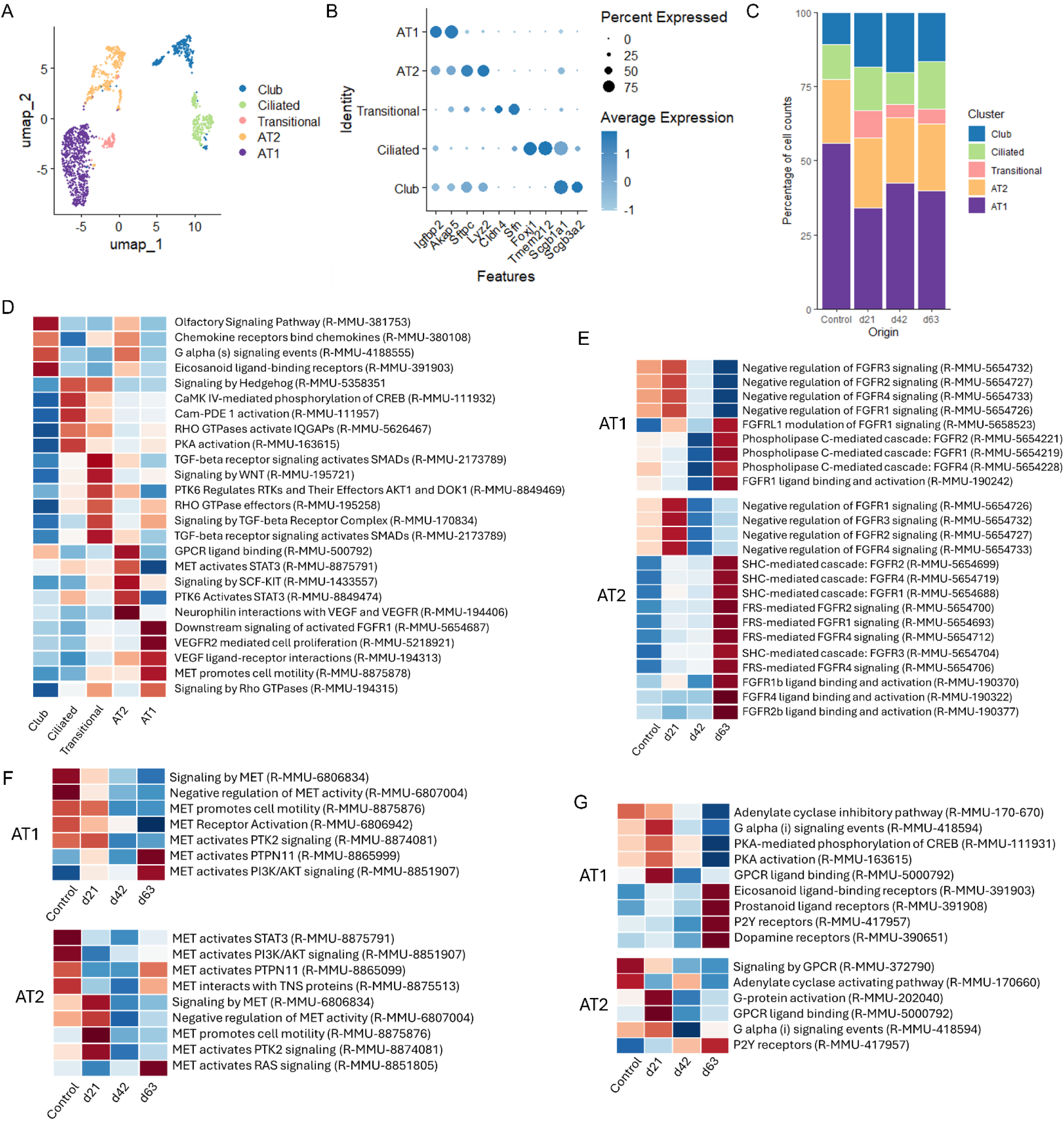
Lung epithelial cells are enriched for pro-regenerative and pro-resolution pathways during spontaneous fibrosis resolution. (A) UMAP of epithelial cell subclusters. (B) Dot plot of representative markers for each epithelial subset. (C) Cluster graph displaying longitudinal cell type composition of epithelial subtypes. (D) GSEA of top enriched pathways among lung epithelial cells. E-G: Longitudinal GSEA of FGF (E), HGF/MET (F), and cAMP signaling and related pathways in AT1 and AT2 cells.

Globally, GSEA identified enrichment of the pro-fibrotic TGFβ pathway (transitional cells) and hedgehog signaling pathway (transitional and ciliated) (Figure 5D). Activation of the putative anti-fibrotic HGF/MET pathway was found to be enriched in AT2 cells and, to a lesser degree, in transitional and ciliated cells. Enrichment of cAMP-related pathways in AT2 and club cells was demonstrated as indicated by enrichment of the stimulatory G-protein (Gαs) that activates adenylyl cyclase and generates intracellular cAMP. Finally, AT1 cells were particularly enriched for activation of the mitogen receptor FGFR1 – a receptor family associated with alveolar regeneration and repair (50, 51).

Longitudinally, GSEA enrichment of profibrotic pathways included enrichment of hedgehog signaling, WNT, and GLI3-related NOTCH signaling during peak fibrosis in AT1 cells and WNT in AT2 cells (Supplemental Fig 5C and D). Mitogen enrichment, specifically FGFR signaling, was found to be negatively regulated at peak fibrosis with a switch towards increased FGFR 1 - 4 signaling by day 63 (Figure 5E). Similar to alveolar and adventitial fibroblasts, AT1 and AT2 cells exhibited negative regulation of HGF/MET activity during peak fibrosis and its interaction with PTK2/FAK with a switch towards downstream effectors such as RAS, PTPN11, and interaction with tenascin proteins by day 63 (Figure 5F). Finally, pathways that modulate intracellular cAMP switched from an inhibitory tone on cAMP generation via Gαi at peak fibrosis to a loss of Gαi enrichment in favor of Gαs-coupled GPCRs by day 63 of resolution (Figure 5G). This pattern of cAMP regulation broadly parallels findings in both macrophages and fibroblasts, namely cAMP activation appears to be inhibited during states of fibrosis and then re-activated during resolution.

### Changes in cellular cross-talk among lung macrophage, fibroblast, and epithelial cells during spontaneous fibrosis resolution

Building on our findings that highlighted increased cellular cross-talk between macrophage populations and fibroblasts (Figure 2B), we sought to pinpoint how the defined macrophage, fibroblast, and epithelial cell subpopulations detailed in Figures 3-5 contribute to the overall cell-cell communication network during fibrosis resolution. To achieve this, we performed CellChat analysis to map signaling interactions among alveolar fibroblasts, adventitial fibroblasts, FAMs, TRAMs, IMs, AT1 cells, and AT2 cells throughout the course of fibrosis resolution. The total number of inferred interactions peaked at day 21 and subsequently declined at day 42 and day 63, indicating increased cell signaling during peak fibrosis that decreases during resolution – though notably not back to baseline levels (Figure 6A). The contribution of each cell subtype to the number of interactions at each time point was determined, demonstrating an increase in interactions among fibroblasts, epithelial cells, and FAMs at peak fibrosis (Figure 6B). The number of interactions among FAMs remained constant throughout resolution while fibroblasts and AT1 displayed a steady reduction over time. AT2 cells were unique in their pattern of interactions that reached a nadir by day 42 with an increase back to baseline at day 63. It is notable that fibroblasts demonstrated the highest number of interactions and exhibited stronger incoming and outgoing signaling activity at each timepoint compared to the other cell types (Supplemental Figure 6A). Further delineation of the specific intercellular pathway interactions at peak fibrosis and during resolution was achieved by comparing the overall information flow for each signaling pathway between day 21 and each resolution timepoint (Supplemental Figure 6B). The net effect of many pathways is difficult to capture by global information flow; therefore, we sought to assess the directionality and relative intensity of signaling among cell types at peak fibrosis and throughout resolution. First, outgoing gene patterns – functionally linked groups of gene programs or pathways – were assigned to every cell subtype at each time point (Supplemental Figure 7, Supplemental Table 1). In saline-treated controls, each cell type exhibited a unique pattern (Supplemental Fig 7A) whereas at peak fibrosis FAMs and TRAMs shared a signaling pattern. At day 42, FAMs established a unique pattern not shared with other cell types and then adopted a pattern similar to IMs by day 63 (Supplemental Figure 7C, D & Supplemental Table 2).

**Figure 6:**
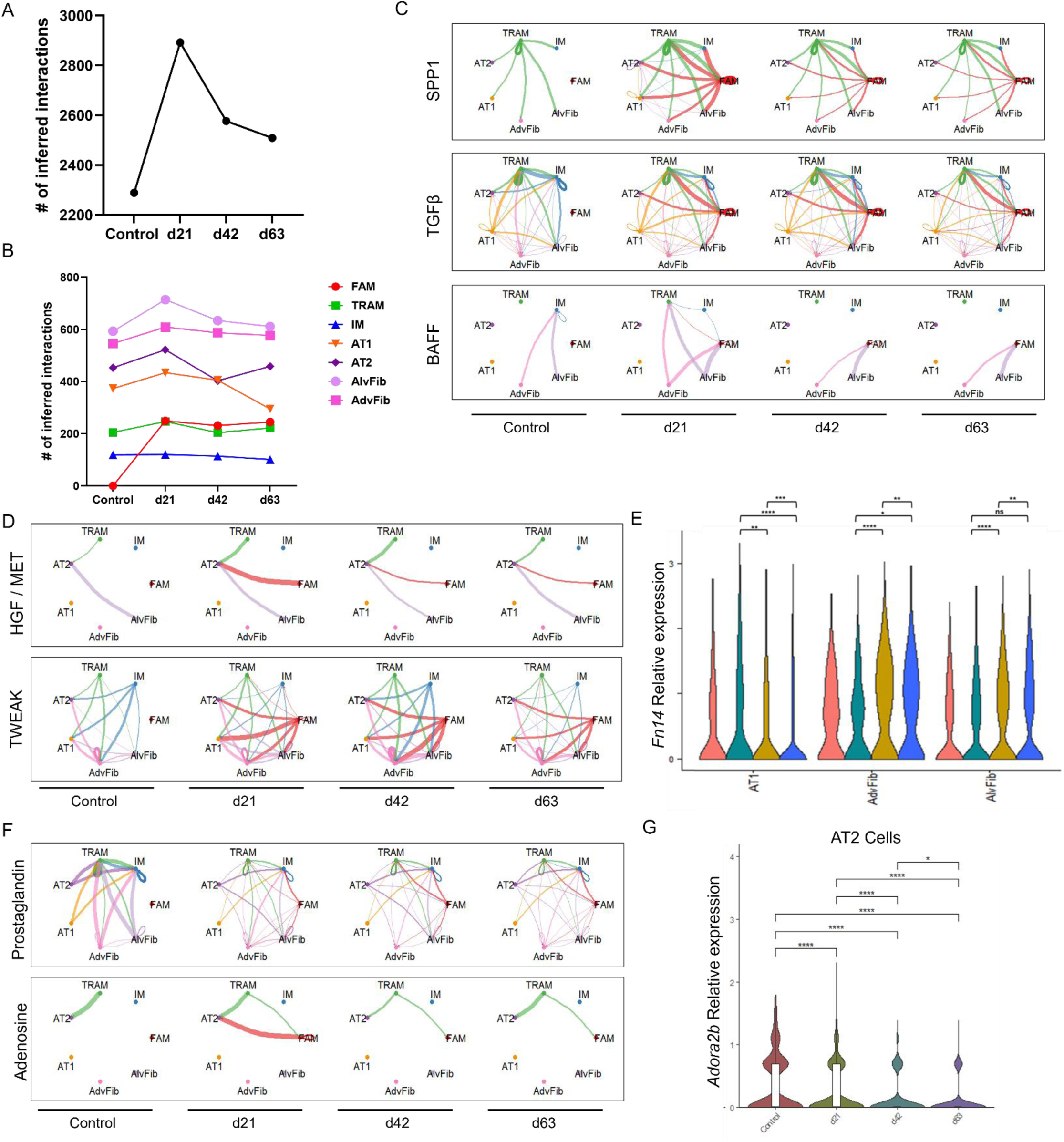
Changes in cellular cross-talk among lung fibroblast, macrophage, and epithelial cells during spontaneous fibrosis resolution. A & B: Longitudinal quantification of the total (A) and cell-specific (B) number of inferred interactions among alveolar fibroblasts, adventitial fibroblasts, FAMs, TRAMs, IMs, AT1 cells, and AT2 cells. C, D, and F: Circos plots depicting longitudinal cellular cross-talk of the profibrotic pathways SPP1, TGFβ, and BAFF (C), anti-fibrotic HGF/MET and TWEAK (D), and cAMP-related Prostaglandin and Adenosine (F) among the indicated cell subtypes. (E) Relative longitudinal expression of the TWEAK receptor *Fn14* transcript in AT1, adventitial fibroblasts, and alveolar fibroblasts. (G) Relative longitudinal expression of the Gαs-coupled *Adora2b* transcript in AT2 cells. Circos plots: line thickness is proportional to signaling strength, line color indicates sender cell.

Circos plots demonstrating relative signaling strength and directionality between cell types were generated for select pro-fibrotic or pro-resolution pathways predicted to be relevant during spontaneous fibrosis resolution by GSEA (Figure 3-5). Among profibrotic programs, SPP1 was produced by TRAMs and signaled to each other cell type present in control mice. FAMs contributed substantially as a source cell of SPP1 during peak fibrosis with minor contributions from AT1, AT2, and IMs. Interestingly, although SPP1 signaling declined in intensity and no longer exhibited outgoing signaling from non-immune cells during resolution, FAMs continued to produce SPP1 at day 63 (Figure 6C, top row of panels). A similar pattern was found with the TGFβ pathway in which FAMs exhibited substantial outgoing signal during peak fibrosis that remained at day 63 (Figure 6C, middle row of panels). Another profibrotic pathway, BAFF (B-cell activating factor) (52), was found to display more specific intercellular interactions. At baseline, IMs appear to receive BAFF signals from alveolar and adventitial fibroblasts with a shift in the recipient cell type to TRAMs and FAMs during peak fibrosis. Furthermore, FAMs become the exclusive recipient of BAFF signals by day 42 with a reduction in signaling intensity from fibroblasts by day 63 (Figure 6C, bottom row of panels).

The persistence of profibrotic SPP1 and TGFβ signaling among cell types through day 63 suggests that the appearance of pro-resolution pathways – and not simply a decline in pro-fibrotic pathways – may be necessary to achieve spontaneous resolution. We therefore assessed anti-fibrotic molecular pathways which were predicted to be enriched by GSEA. Among them, the pro-resolution MET/HGF axis was found to be produced by alveolar fibroblasts and TRAMs at baseline with FAMs contributing during peak fibrosis and throughout resolution, albeit with lower intensity. Interestingly, AT2 cells were the only cell predicted to receive MET/HGF signals (Figure 6D, top). TWEAK signaling at baseline demonstrated cross-talk among immune cells, fibroblasts, and epithelial cells with a substantial increase in outgoing signaling from FAMs to fibroblasts and epithelial cells on day 42 (Figure 6D, bottom). The strength of TWEAK signaling in recipient cells is dependent upon the expression of its sole receptor, FN14. We therefore assessed the relative expression of FN14 transcript and found that its expression increased at day 42 and day 63 while declining in AT1 cells by day 63 (Figure 6E).

Pathways that signal through cAMP were also assessed. Prostaglandin signaling was robust among most cell types in control lungs with immune, AT1/AT2, and fibroblasts serving as cellular sources. During peak fibrosis and resolution, outgoing prostaglandin signaling from AT2 and IM cells appeared to expand to now include fibroblasts as recipient cells (Figure 6F, top). Adenosine, which signals via the GPCRs Adora 1, 2a, 2b, and 3 demonstrated baseline TRAM-to-AT2 communication which expanded to include FAMs at day 21 with a return to baseline at day 42 and day 63 (Figure 6F, bottom). Given that AT2 cells are predicted to be the main recipient cell of adenosine, we measured expression levels of Adora genes and found that only Adora 2b, a Gαs-coupled receptor, was appreciably expressed with its highest expression at baseline and peak fibrosis (Figure 6G).

## DISCUSSION

Therapeutic drug targets for fibrotic lung disease have historically been selected primarily for their ability to inhibit putative drivers of fibrosis. This strategy reflects the fact that a vast majority of fibrosis research has focused solely on the cellular and molecular determinants of fibrogenesis. By contrast, an alternative research paradigm seeking to explicate the endogenous mechanisms that underly fibrosis resolution may identify essential processes absent during persistent fibrosis that are candidates for therapeutic restoration (1). In particular, the cellular subtypes, signaling pathways, and cross-talk that might enable and/or promote fibrosis resolution as observed in the single-dose murine bleo model have not been characterized. In this study we performed scRNA-seq to determine the longitudinal cellular landscape, pathway enrichment, and cellular cross-talk operative during spontaneous fibrosis resolution in this model as it occurs in young normal healthy mice. Our findings highlight conspicuous shifts in FAM numbers and signaling as well as dynamic shifts in a recently characterized matrix-secreting fibroblast subtype (*Csmd1*+). Moreover, while signaling of many pro-fibrotic pathways were found to diminish by days 42 and 63 of resolution, our findings also suggest that anti-fibrotic MET/HGF, cAMP, and TWEAK signaling and cellular cross-talk is enriched during resolution.

A hallmark of lung fibrosis is the persistence of aberrant cellular phenotypes that contribute to its defining pathobiological features, including alveolar damage, dysregulated cellular integrity and survival, and excessive matrix deposition (1, 53). We therefore predicted that shifts in cellular subtypes among immune, mesenchymal, and epithelial cells would be evident throughout resolution. To our surprise, we did not observe substantial changes in the proportions of fibroblast or epithelial subtypes evident in UMAPs. Though a small transitional epithelial cell cluster was observed at day 21, this is likely to be a remnant population that peaked earlier during the fibrotic response (49) with gradual decline throughout resolution. It will be important for future studies to determine the possible role of transitional epithelial cells in models of persistent fibrosis. Among mesenchymal cells, we did identify a small *Csmd1*+ fibroblast population present within alveolar, adventitial, and peribronchial subsets recently characterized to be pro-fibrotic (19). Although the wider significance of this fibroblast population remains to be further characterized in future studies, we found that it expressed high levels of pro-fibrotic genes (SPP1 and LTBP2) and that it declined by day 42 and through day 63 (Fig. 4F). The lack of substantial *Cthrc1* expression in our model is notable and possibly explained by the lower dose of bleo (1.0 U/kg) we employed compared to other studies (54) as well as the lack of fibroblast enrichment.

The most conspicuous longitudinal cellular change in our study was among FAMs. Essentially absent in control lungs, FAMs expanded and peaked at day 21 following bleo, gradually declining throughout resolution (Figure 3C, Supplemental Figure 3A). Similar macrophage cell populations expressing fibrosis-associated genes as well as IM and TRAM markers have been identified and characterized during bleo-induced inflammatory and fibrotic phases by other groups (10, 15, 38, 40). It has been proposed that these cells may be derived from IMs and ultimately transition into AMs (10, 39). The recent study by Lin *et al* (38) advances this notion by demonstrating the emergence of a pro-fibrotic “Mac0” population by day 7 post-bleo originating from recruited monocytes and persisting through day 28. Furthermore, the study demonstrates that recruited monocyte-derived macrophages undergo an IM-to-AM transition via a Mac0 transient state. In our study, FAMs displayed a close UMAP alignment to IMs and TRAMs similar that of the Mac0 population (38). Additionally, the relative proportions of FAMs on day 21 are consistent with the Mac0 trends between days 14 and 28. These prominent parallels suggest that FAMs and Mac0s may indeed represent the same cell population. Although the paucity of FAMs in untreated mouse lungs suggests that they represent a recruited population, our study did not assess earlier time points following bleo administration (days 3, 7, and 14) or utilize lineage tracing to determine whether FAMs are indeed recruited. Furthermore, we acknowledge that the pro-fibrotic phenotype of these macrophages is associative and has not been experimentally demonstrated.

Our finding that epithelial, mesenchymal, and immune cell subtype proportions (except for FAMs) remained essentially constant during resolution strongly suggests that more subtle changes – inconspicuous by UMAP – might be responsible for fibrosis resolution and a trajectory toward re-establishing lung homeostasis. Furthermore, as lung homeostasis and repair reflect the dynamic communication among immune, mesenchymal, and epithelial cells (55–57), shifts in cellular crosstalk are likely to be an important part of the resolution process. Indeed, our single-cell analysis confirmed a rich interplay of bidirectional signaling among these cell types during fibrosis and its spontaneous resolution (Figure 2B). GSEA revealed a decline in pro-fibrotic signaling over time in immune, mesenchymal, and epithelial cell subtypes, including NOTCH (fibroblasts and FAMs), Hedgehog (fibroblasts, AT1), and WNT signaling (fibroblasts, TRAMs, FAMs) (Figure 3H and 3J, Supplemental Figure 3C, 4D, 5C & D). Additionally, a reduction in signaling strength was observed in SPP1, TGFβ, and BAFF signaling (Figure 6C), consistent with what would be expected in a resolving environment. However, these signaling strength or cross-talk patterns (in the case of BAFF) did not return all the way to baseline. Other pathways such as MIF and PDGF did not appear to appreciably diminish, instead maintaining a complex network of cellular cross-talk (Supplemental Figure 8). The pro-fibrotic actions of MIF depend on its ligation of CD74 in Cx3cr1 expressing lung macrophages (32). In our study, FAMs and IMs expressed the highest transcript levels of Cx3cr1 (Figure 3B) and it is thus notable that only the canonically pro-fibrotic FAMs exhibited a decline in CD74 expression during resolution, though not to baseline levels (Supplemental Figure 9). Such persistent enrichment of some pro-fibrotic signaling pathways may reflect the fact that complete restoration of cellular homeostasis requires longer than 63 days or that, despite evidence of histopathologic resolution, there is a kind of cellular priming that persists following injury. The latter explanation is consistent with the observation that repetitive injury models lead to persistent fibrosis (58). The pathways GDF and GAS – which can contribute to either pro-fibrotic or pro-resolution signaling (59–61) – were also highly enriched in control and bleo-treated lungs at each time point (Supplemental Figure 8). It is possible that ongoing pro-fibrotic signaling at days 42 and days 63 must be counterbalanced by concomitant upregulation or switching to pro-resolution pathways.

Indeed, longitudinal analysis revealed enrichment of several putative pro-resolution pathways over time. The HGF/MET axis was found to be enriched in AT2 cells where it has been demonstrated to favor epithelial restitution (62, 63) as well as in fibroblasts where it functions to oppose TGFβ signaling and sensitize them to apoptosis (64, 65). The negative regulation of MET on day 21 with a switch to its activation of downstream targets (such as PTPN11, RAS, TNS proteins) by day 63 is consistent with such pro-resolving functions. The enrichment of MET-mediated activation of PTK2/FAK at day 21 in fibroblasts, which is predicted to confer pro-fibrotic properties (66, 67), diminishes by day 42. Another signaling switch observed during resolution involves FGF signaling within alveolar epithelial cells. The inhibition of FGFRs at day 21 stands in contrast to their activation by day 63 in AT1 and AT2 cells (Figure 5E). FGFR2 signaling in AT2 cells is known to favor their maintenance and recovery after injury (28) and to regulate the balance of AT2-to-AT1 differentiation (51, 68). As FGF2 is known to expand fibroblast populations through proliferation and promote their survival (69, 70), nintedanib (an inhibitor of FGFR2, VEGFR, and PDGFR) is approved to treat IPF and progressive pulmonary fibrosis. However, given the pro-resolution effects of FGFR2 signaling within alveolar epithelial cells, one might speculate that nintedanib could limit its own efficacy by hampering AT2 maintenance.

The significance of TWEAK enrichment at peak fibrosis and throughout resolution (Figure 6D) depends upon its interaction with its receptor Fn14 (71). Furthermore, although TWEAK-Fn14 signaling has been reported to exhibit anti-fibrotic effects in the lung (34), it has similarly been found to promote fibrosis in the kidney (72). This dichotomy may be attributed to fibroblast deactivation and recruitment of pro-resolution macrophages in the lung (34) as opposed to fibroblast expansion and inflammation in the kidney (73). Sustained epithelial TWEAK-Fn14 signaling could also account for its profibrotic actions in the kidney and other organs (74). Assessing the longitudinal expression of *Fn14* in epithelial cells and fibroblasts in our model, we found that *Fn14* declined substantially by day 63 in AT1 cells and increased its expression in fibroblasts throughout resolution (Figure 6E). Dynamic and complementary expression patterns of Fn14 in AT1 cells and fibroblasts during spontaneous resolution thus appear to be consistent with the previously described anti-fibrotic effects of TWEAK-Fn14 in the lung.

An important and consistent finding from longitudinal GSEA is the temporal regulation of intracellular cAMP signaling. Specifically, inhibition of adenylyl cyclase via Gαi was found to occur at peak fibrosis with a switch to cAMP pathway activation evidenced by Gαs-coupled GPCR or PKA activation of CREB enrichment by day 63 (Figure 3I, 4I, 5G). This pattern is consistent with the putative anti-fibrotic actions of cAMP within immune, mesenchymal, and epithelial cell populations in the lung (75). Though these observations suggest a potential endogenous role for cAMP during resolution, the operative GPCRs responsible for generating intracellular cAMP within each of the pivotal cell types remain unclear. Neither receptors for prostaglandin family members nor adenosine – which can each activate or inhibit adenylyl cyclase through various GPCRs – revealed concrete patterns consistent with the timely activation of cAMP following peak fibrosis observed in each cell type by GSEA. One exception may be adenosine signaling in AT2 cells which primarily expressed the Gαs-coupled *Adora2b* transcript (Figure 6G). It is conceivable that early endogenous pro-resolution signaling may overlap with peak fibrosis. Consistent with the reported ability of cAMP via prostaglandin signaling to increase fibroblast expression of the death receptor Fas (76), both alveolar and adventitial fibroblasts showed enrichment in FasL signaling – a crucial step for spontaneous resolution (6). The potential contributions of cAMP signaling to spontaneous fibrosis resolution in our model are especially notable considering that the recently approved IPF drug nerandomilast functions to inhibit the breakdown of cAMP (77) as well as the recent phase 3 success of stimulating cAMP generation through inhaled treprostinil in IPF (78).

This study has several limitations. First, single-cell sequencing, by its nature, is associative and hypothesis-generating, as mRNA transcripts do not fully correlate with protein expression and cannot precisely predict cellular phenotype/function. It is therefore important that pathways and cell-cell interactions that plausibly impact fibrosis resolution be confirmed in the appropriate experimental context. Next, our use of a relatively low dose of bleo (1.0 U/kg) to ensure timely resolution may, for example, account for the lack of *Cthrc1*+ fibroblasts observed during peak fibrosis and thus does not fully represent the extent of lung fibrosis in human IPF. We acknowledge that our findings will need to be independently reproduced with a larger number of mice. Finally, the significance of the potentially pro-resolving cellular transitions, signaling pathways, and cross-talk identified herein would be substantially strengthened if these were shown to be impaired or deficient in experimental models of non-resolving and persistent fibrosis. Overall, our findings suggest a rich milieu of cellular cross-talk in the lung as well as dynamic enrichment of pro-fibrotic and pro-resolution pathways during peak fibrosis and throughout fibrosis resolution. Additional interrogation of these pathways and patterns in the future may further illuminate our understanding of fibrosis outcomes and their therapeutic modulation.

## Acknowledgments

This work was funded by NIH grants R35 HL144979 (to MPG), K08 HL163178 (to SMF), R35 HL160770 (to RLZ), R01 HL175555 (to TB), Hevolution HF-AGE award (C.A.A.), National Science Foundation CAREER award (2045977), the Department of Defense and Congressionally Directed Medical Research Program W81XWH2010336 and W81XWH2110491 (C.A.A.), Defense Advanced Research Projects Agency (DARPA) “BETR” award D20AC0002 (C.A.A.) awarded by the U.S. Department of the Interior (DOI), Interior Business Center. The content is solely the responsibility of the authors and does not necessarily represent the official views of the National Institutes of Health or National Science Foundation, the position or the policy of the Government, and no official endorsement should be inferred. All Figure schematics were created with BioRender (BioRender.com).

## Author contributions

JS, NMW, MPG, and SMF designed the experiments. Experiments were performed by JS, NMW, and SMF. Bioinformatics were performed by JS, VW, and YL. Data were analyzed by JS, VW, SG, CA, MPG, and SMF. Intellectual contributions were provided by TB and RZ. The manuscript was written by JS and SMF. All authors reviewed the manuscript.

## SUPPLEMENTAL FIGURES

**Supplemental Figure 1:**
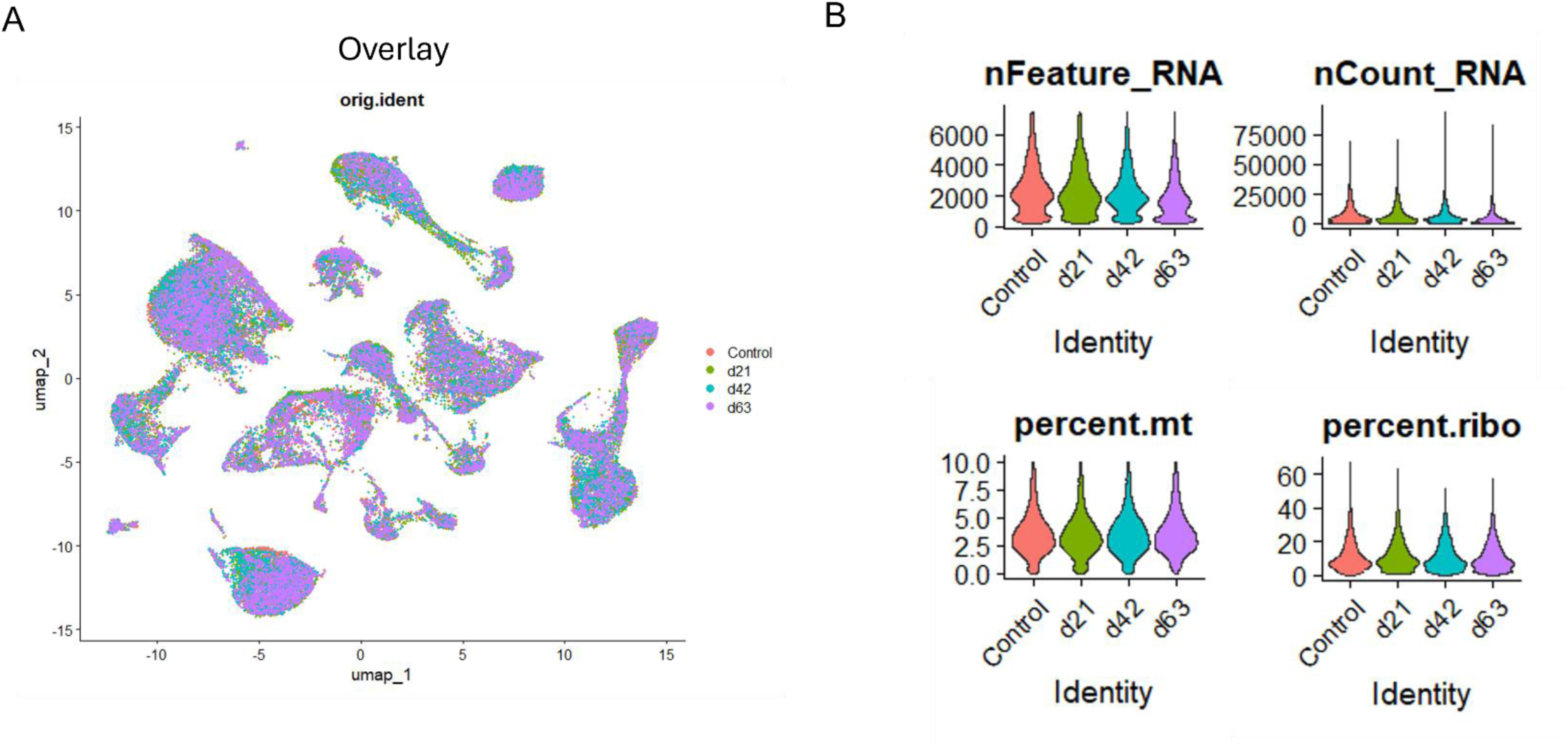
(A) Superimposed Uniform Manifold Approximation and Projection (UMAP) of all conditions and time points. (B) Standard quality control thresholds for unique genes/cell (upper left), unique molecular identifiers/cell (upper right), mitochondrial percentage (lower left), and ribosomal percentage (lower right).

**Supplemental Figure 2:**
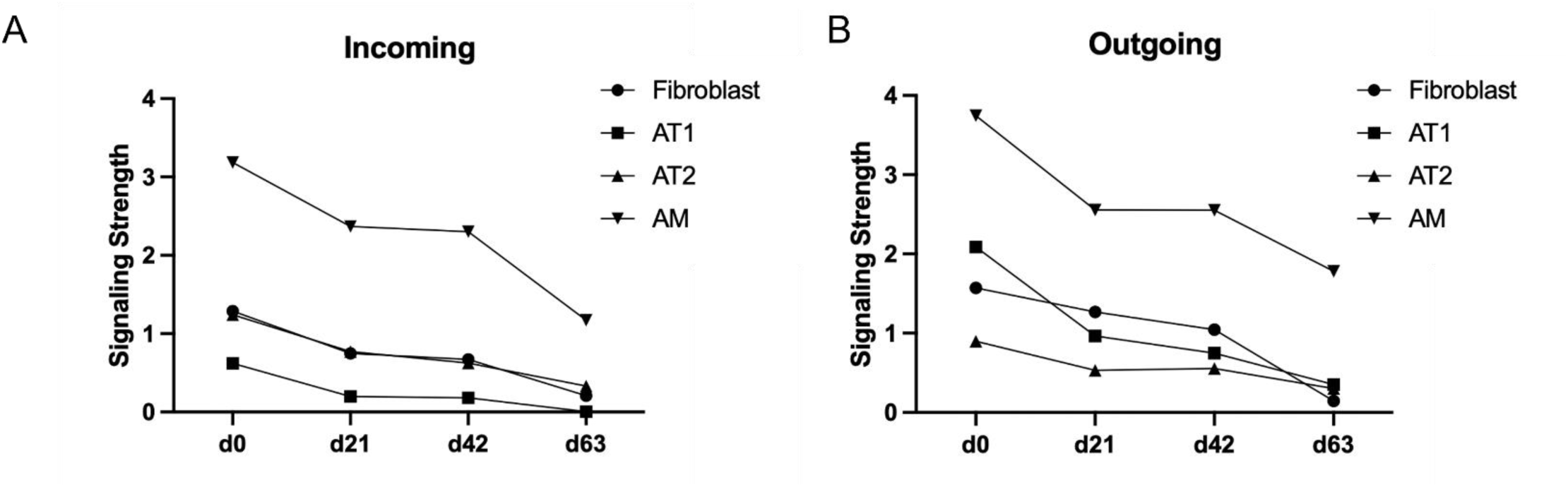
Relative incoming (A) and outgoing (B) signaling strength of fibroblast, AT1, AT2, and AMs quantified longitudinally.

**Supplemental Figure 3:**
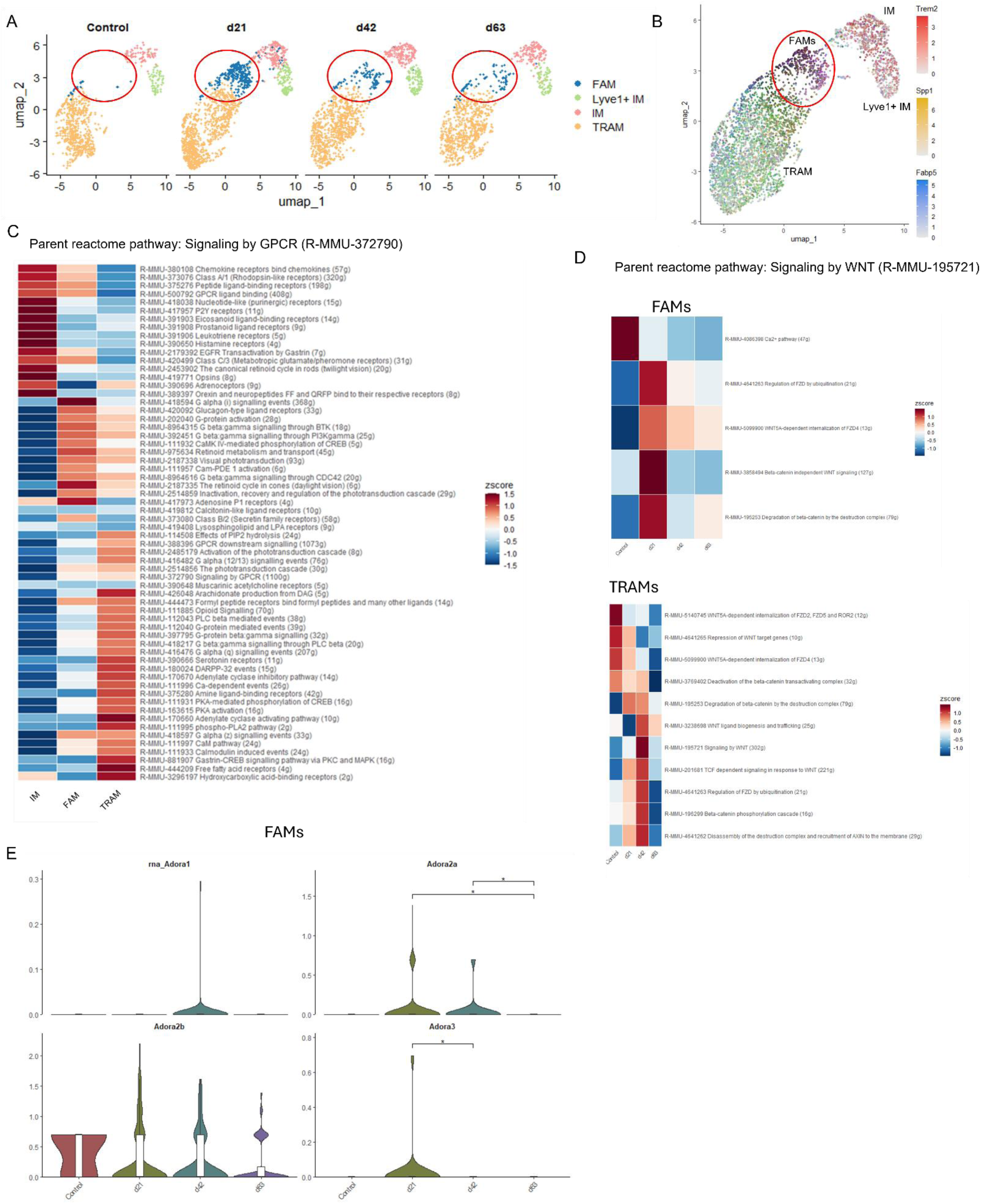
(A) Longitudinal UMAP of lung macrophage subsets during resolution. Red circle encompasses FAM cell population. (B) Feature plot of lung macrophage subsets displaying relative expression of Trem2, Spp1, and Fabp5. (C) GSEA of macrophage subsets for “signaling by GPCR” parent reactome pathway. (D) GSEA of “signaling by WNT” parent reactome pathway in FAMs (top) and TRAMs (bottom). (E) Violin plots displaying relative transcript expression of Adora 1, 2a, 2b, and 3 longitudinally.

**Supplemental Figure 4:**
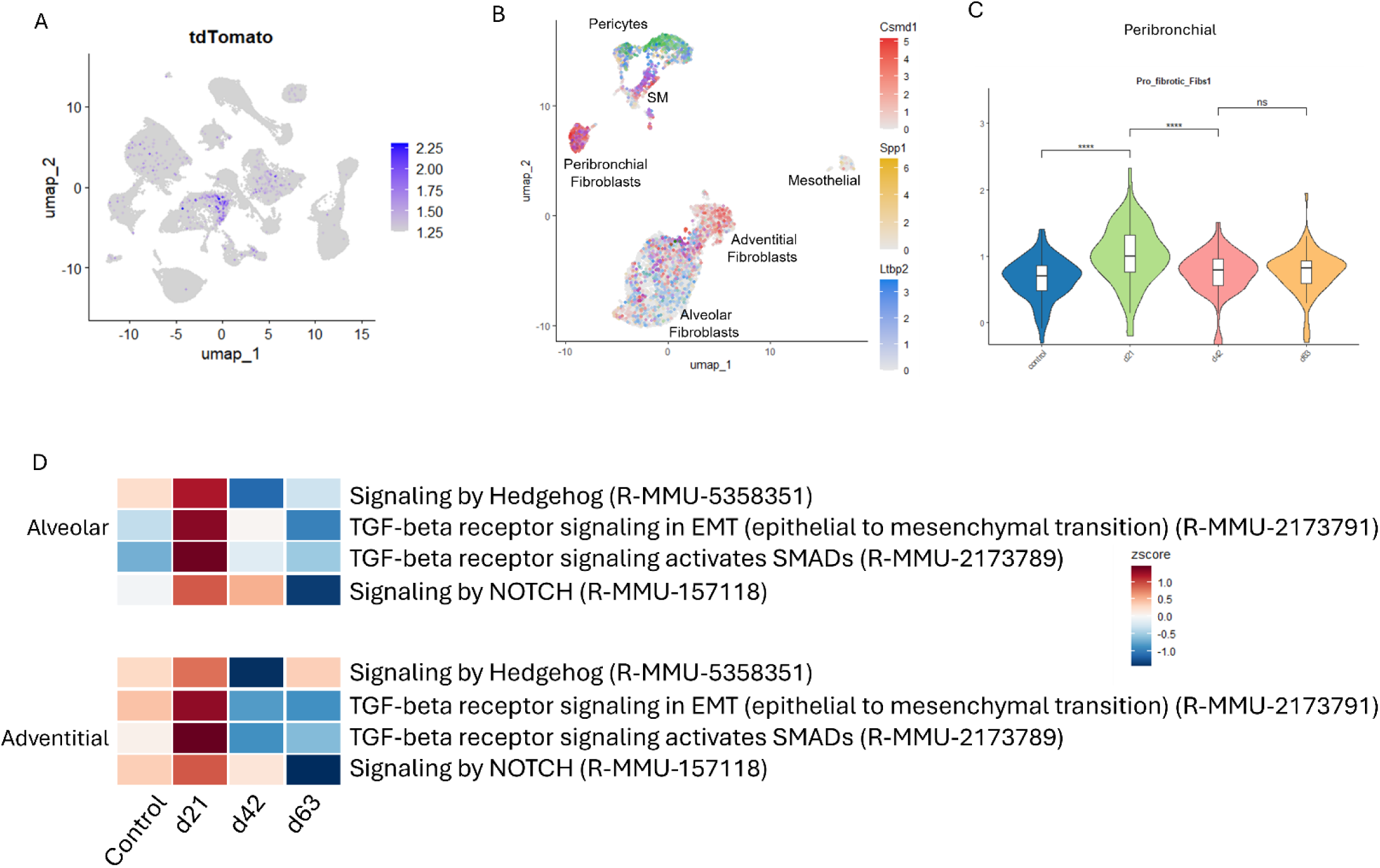
(A) Feature plot of tdTomato expression among all lung cell types. (B) Feature plot of lung mesenchymal subsets displaying relative expression of Csmd1, Spp1, and Ltbp2. (C) Violin plot displaying relative presence of “ECM secreting” fibroblast subtype longitudinally within peribronchial cells defined by the feature plot in B. (D) Longitudinal GSEA of selected pathways within alveolar and adventitial fibroblast subtypes.

**Supplemental Figure 5:**
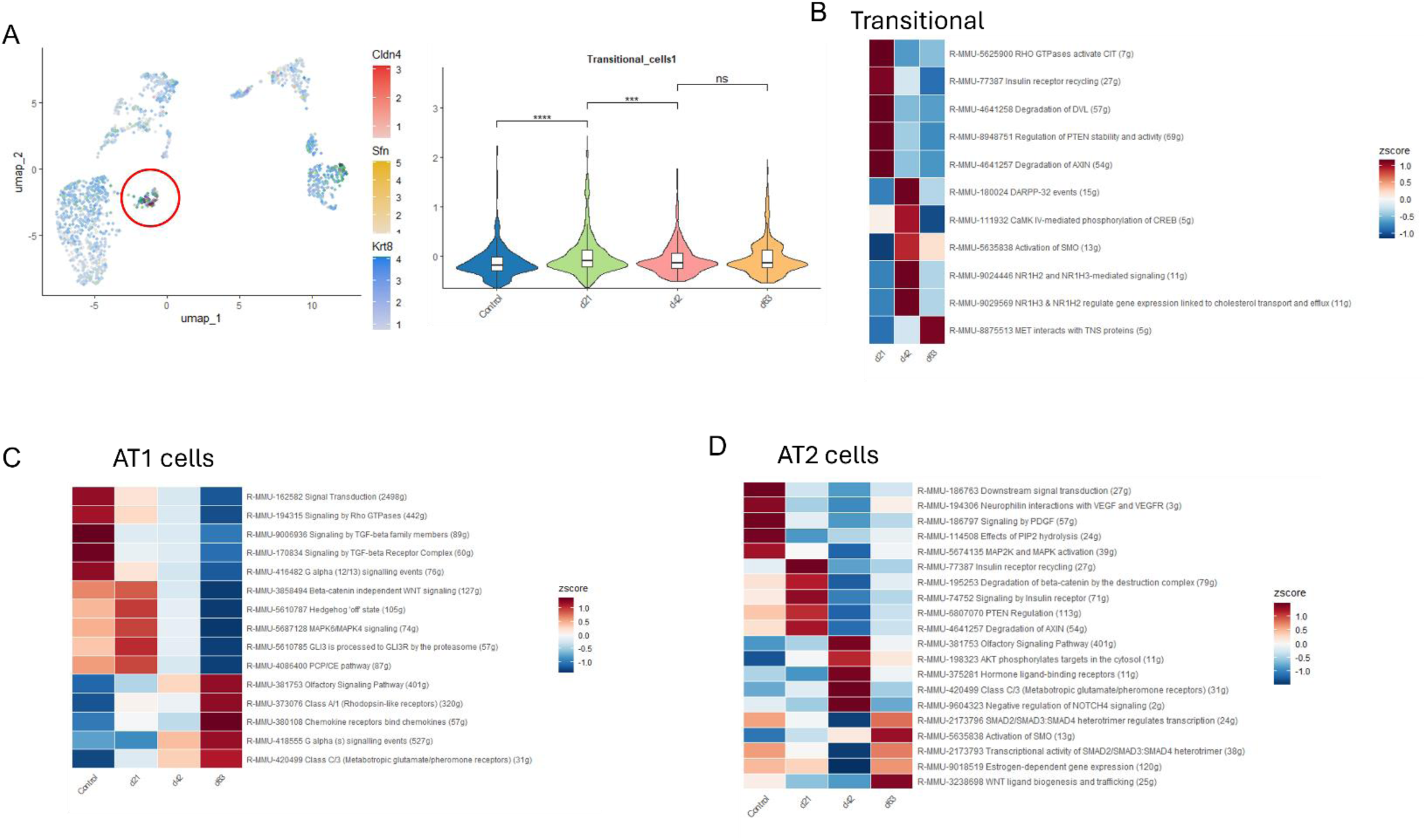
(A) Feature plot of Cldn4, Sfn, and KRT8 expression to identify transitional epithelial cell population (red circle). Longitudinal GSEA of top pathways in transitional (B), AT1 cells (C), and AT2 cells (D).

**Supplemental Figure 6:**
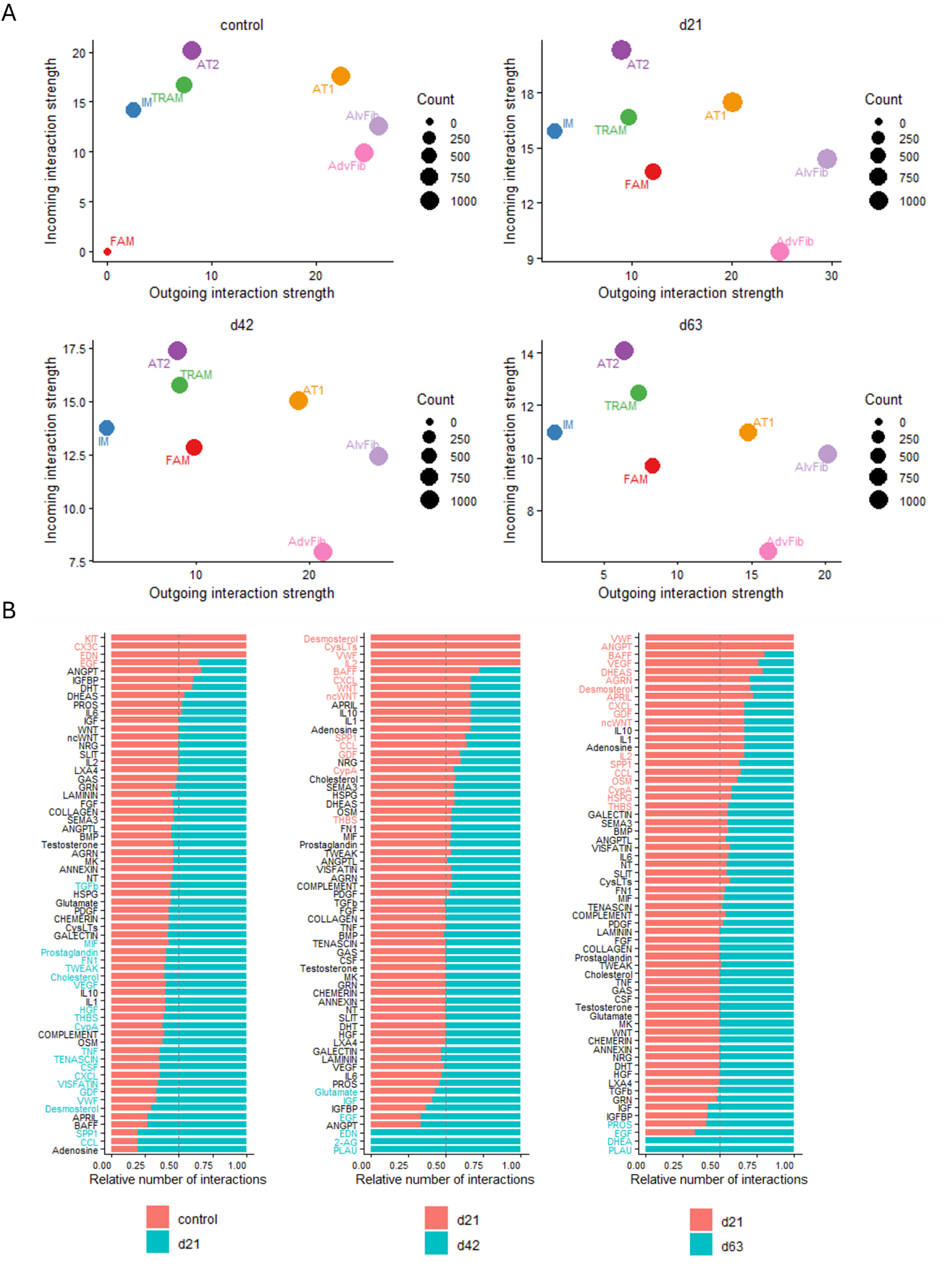
(A) Dot plot depicting relative incoming and outgoing interaction strength the indicated cell types in control and bleomycin treated mice at each time point. (B) Waterfall plots displaying relative enrichment of the indicated pathways. Each plot compares a condition or timepoint (control – left, day 42 bleo – middle, day 63 bleo – right) to day 21 bleo-treated mice.

**Supplemental Figure 7:**
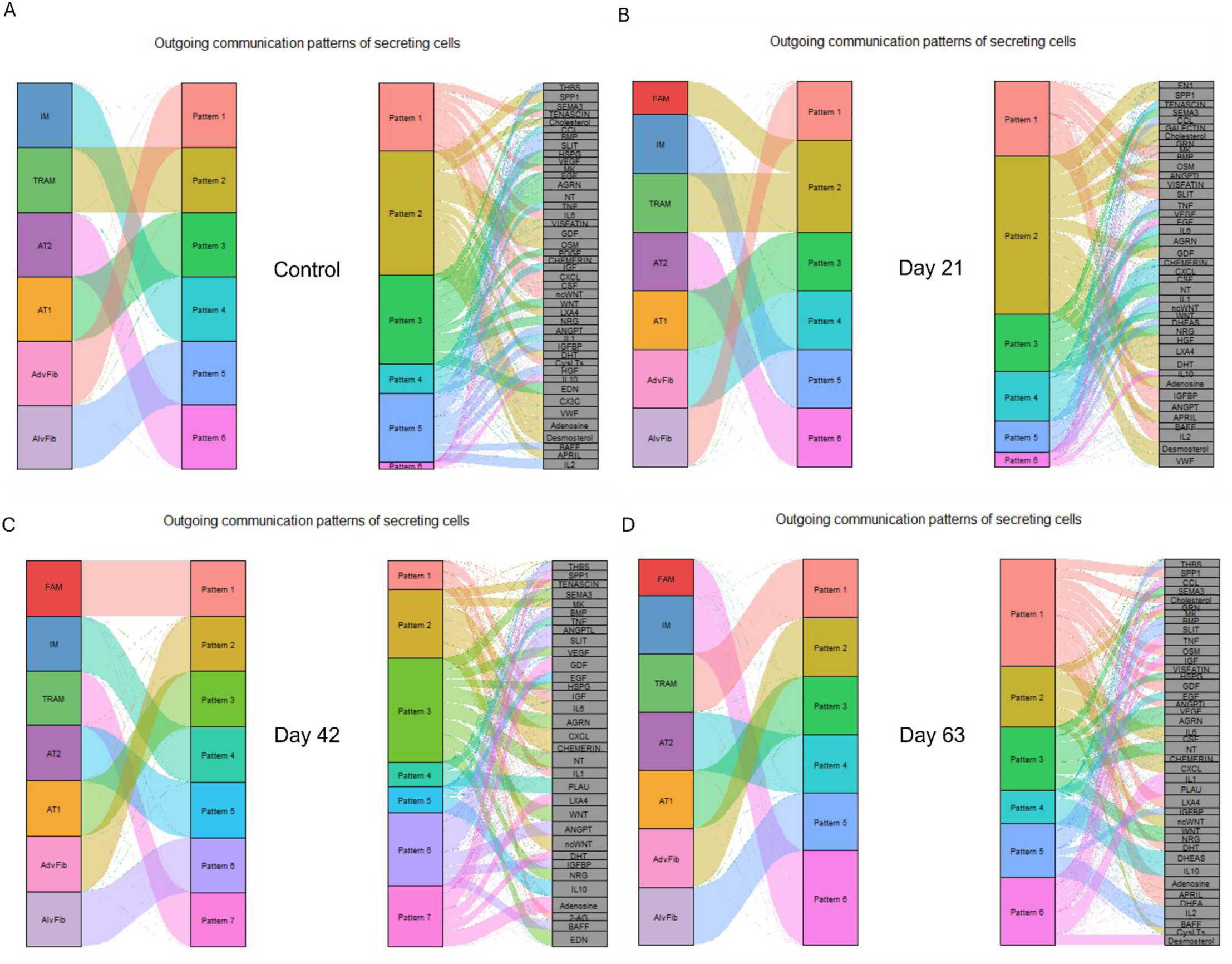
Alluvial plots of outgoing gene patterns for the indicated cell types (left) and genes that define each pattern (right) in control (A), day 21 bleo (B), day 42 bleo (C), and day 63 bleo (D).

**Supplemental Figure 8:**
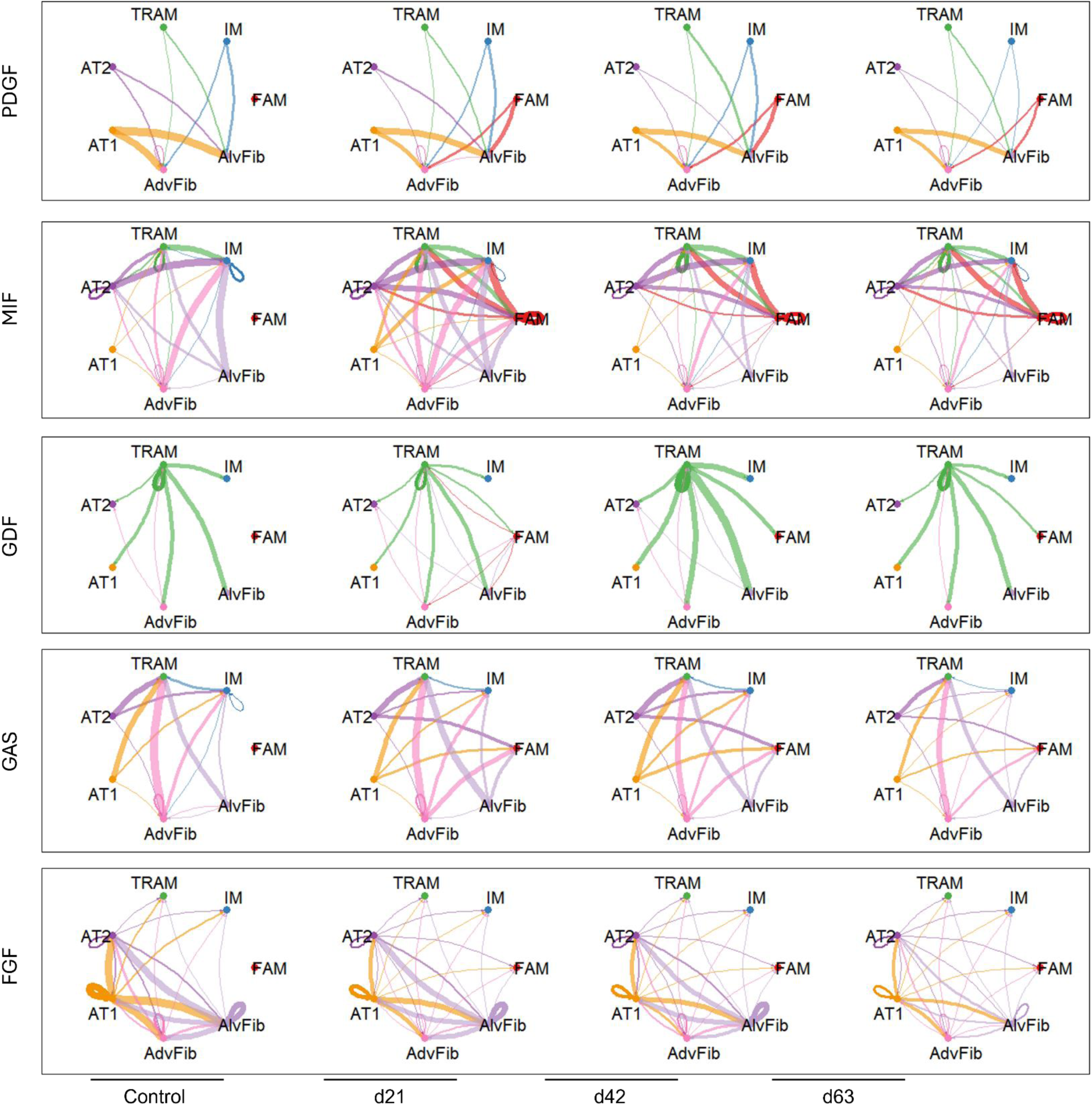
Circos plots displaying longitudinal cellular cross-talk of the indicated pathways. Line thickness is proportional to signaling strength, line color indicates sender cell.

**Supplemental Figure 9:**
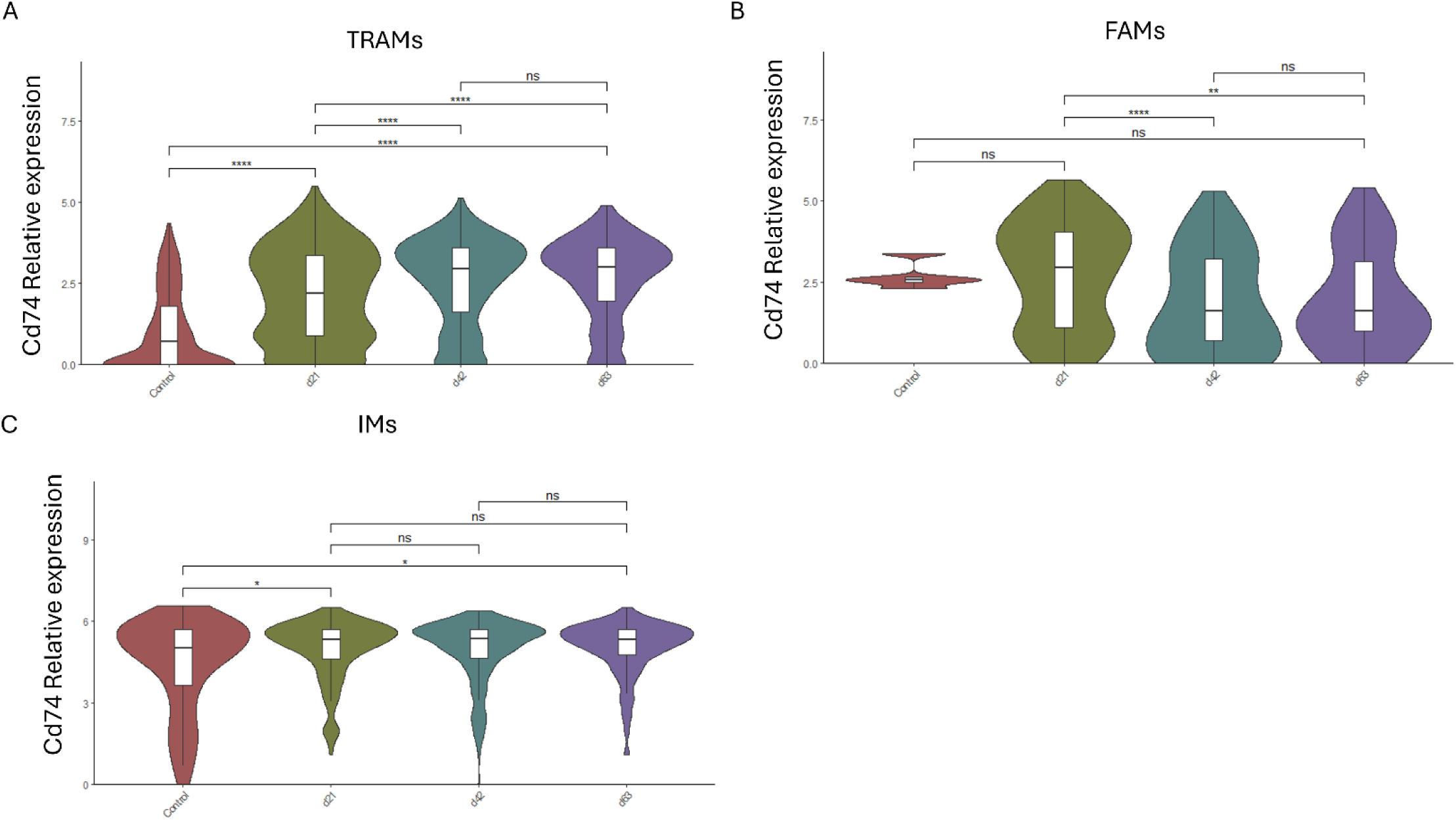
Violin plots displaying relative longitudinal *Cd74* transcript expression in TRAMs (A), FAMs (B), and IMs (C).

**Supplemental Table 1:**
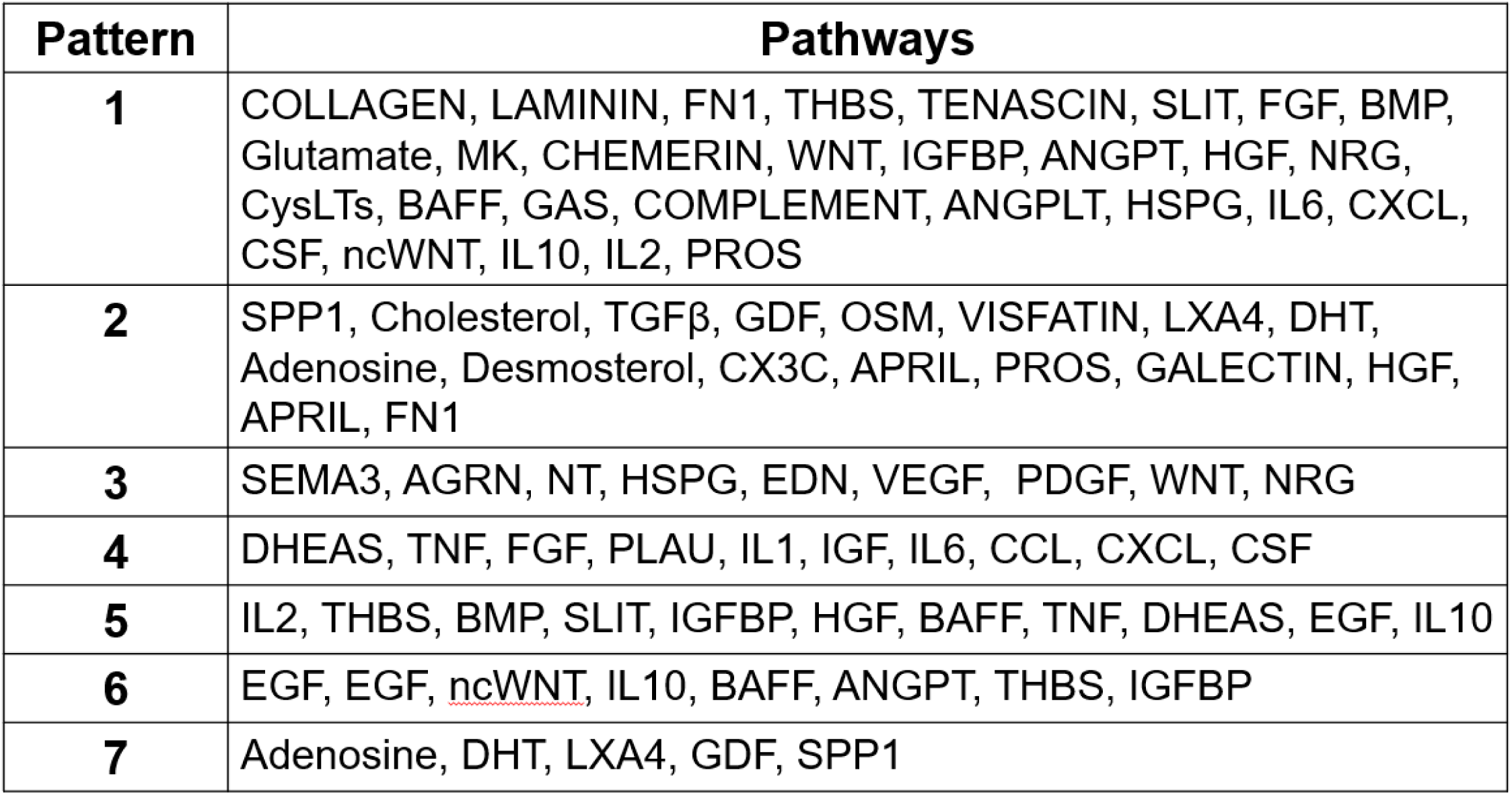
List of top enriched genes that define each outgoing gene pattern derived from alluvial plots in Supplemental Figure 7.

**Supplemental Table 2:**
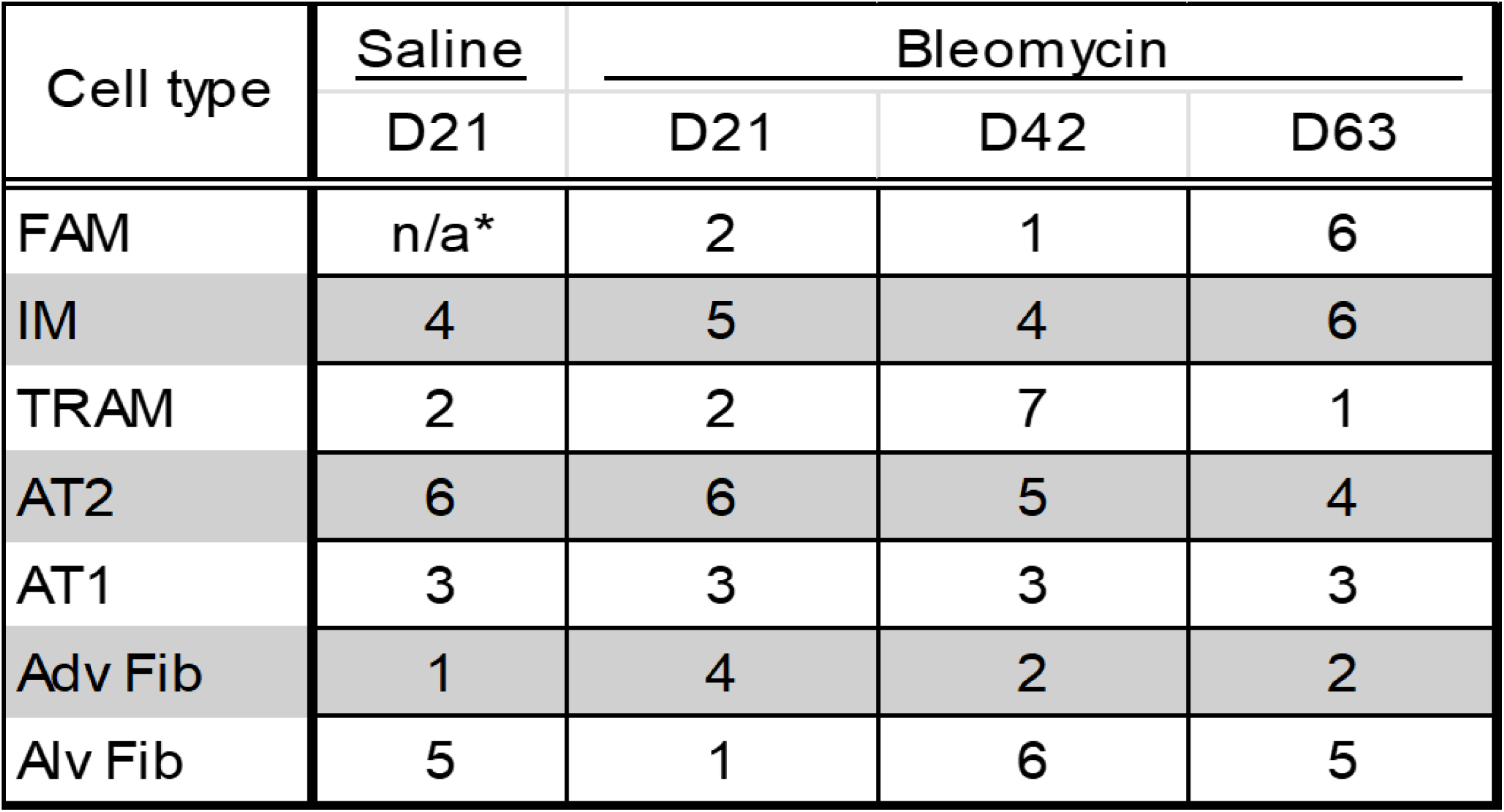
Longitudinal gene pattern identity for each cell type from alluvial plots in Supplemental Figure 7.

| Cell type | Saline | Bleomycin |  |  |
| --- | --- | --- | --- | --- |
|  | D21 | D21 | D42 | D63 |
| FAM | n/a* | 2 | 1 | 6 |
| IM | 4 | 5 | 4 | 6 |
| TRAM | 2 | 2 | 7 | 1 |
| AT2 | 6 | 6 | 5 | 4 |
| AT1 | 3 | 3 | 3 | 3 |
| Adv Fib | 1 | 4 | 2 | 2 |
| Alv Fib | 5 | 1 | 6 | 5 |

## REFERENCES

1. Fortier SM, Redente EF, Peters-Golden M. Reimagining Fibrosis Research, Outcomes, and Therapeutics Through the Lens of Resolution. Semin Respir Crit Care Med 2025; 46: 298–310.

2. Glasser SW, Hagood JS, Wong S, Taype CA, Madala SK, Hardie WD. Mechanisms of Lung Fibrosis Resolution. Am J Pathol 2016; 186: 1066–1077.

3. B BM, Lawson WE, Oury TD, Sisson TH, Raghavendran K, Hogaboam CM. Animal models of fibrotic lung disease. Am J Respir Cell Mol Biol 2013; 49: 167–179.

4. Lawson WE, Polosukhin VV, Stathopoulos GT, Zoia O, Han W, Lane KB, et al. Increased and prolonged pulmonary fibrosis in surfactant protein C-deficient mice following intratracheal bleomycin. Am J Pathol 2005; 167: 1267–1277.

5. Phan SH, Armstrong G, Sulavik MC, Schrier D, Johnson KJ, Ward PA. A comparative study of pulmonary fibrosis induced by bleomycin and an O2 metabolite producing enzyme system. Chest 1983; 83: 44S–45S.

6. Redente EF, Chakraborty S, Sajuthi S, Black BP, Edelman BL, Seibold MA, et al. Loss of Fas signaling in fibroblasts impairs homeostatic fibrosis resolution and promotes persistent pulmonary fibrosis. JCI Insight 2020; 6.

7. Fortier SM, Walker NM, Penke LR, Baas JD, Shen Q, Speth JM, et al. MAPK phosphatase 1 inhibition of p38alpha within lung myofibroblasts is essential for spontaneous fibrosis resolution. J Clin Invest 2024; 134.

8. Pan T, Feng Y, Li Y, Yang Y, Zhou J, Song Y. Exacerbation of pulmonary fibrosis following acute lung injury via activin-A production by recruited alveolar macrophages. Journal of Thoracic Disease 2024; 16: 7709–7728.

9. Penke LR, Speth JM, Huang SK, Fortier SM, Baas J, Peters-Golden M. KLF4 is a therapeutically tractable brake on fibroblast activation that promotes resolution of pulmonary fibrosis. JCI Insight 2022; 7.

10. Aran D, Looney AP, Liu L, Wu E, Fong V, Hsu A, et al. Reference-based analysis of lung single-cell sequencing reveals a transitional profibrotic macrophage. Nature Immunology 2019; 20: 163–172.

11. Jin C, Chen Y, Wang Y, Li J, Liang J, Zheng S, et al. Single-cell RNA sequencing reveals special basal cells and fibroblasts in idiopathic pulmonary fibrosis. Scientific Reports 2024; 14.

12. Mayr CH, Santacruz D, Jarosch S, Bleck M, Dalton J, Mcnabola A, et al. Spatial transcriptomic characterization of pathologic niches in IPF. Science Advances 2024; 10.

13. Reyfman PA, Walter JM, Joshi N, Anekalla KR, McQuattie-Pimentel AC, Chiu S, et al. Single-Cell Transcriptomic Analysis of Human Lung Provides Insights into the Pathobiology of Pulmonary Fibrosis. Am J Respir Crit Care Med 2019; 199: 1517–1536.

14. Vannan A, Lyu R, Williams AL, Negretti NM, Mee ED, Hirsh J, et al. Title (In press).

15. Wang S, Li J, Wu C, Lei Z, Wang T, Huang X, et al. Single-Cell RNA Sequencing Reveals Monocyte-Derived Interstitial Macrophages with a Pro-Fibrotic Phenotype in Bleomycin-Induced Pulmonary Fibrosis. International Journal of Molecular Sciences 2024; 25: 11669.

16. Zhang T, Hou Z, Ding Z, Wang P, Pan X, Li X. Single Cell <SCP>RNA</SCP>-Seq Identifies Cell Subpopulations Contributing to Idiopathic Pulmonary Fibrosis in Humans. Journal of Cellular and Molecular Medicine 2025; 29.

17. Zhou Y, Tong Z, Zhu X, Wu C, Zhou Y, Dong Z. Deciphering the cellular and molecular landscape of pulmonary fibrosis through single-cell sequencing and machine learning. Journal of Translational Medicine 2025; 23.

18. Zhu M, Yi Y, Jiang K, Liang Y, Li L, Zhang F, et al. Single-cell combined with transcriptome sequencing to explore the molecular mechanism of cell communication in idiopathic pulmonary fibrosis. Journal of Cellular and Molecular Medicine 2024; 28.

19. Guo JL, Griffin M, Yoon JK, Lopez DM, Zhu Y, Lu JM, et al. Histological signatures map anti-fibrotic factors in mouse and human lungs. Nature 2025; 641: 993–1004.

20. McGinnis CS, Murrow LM, Gartner ZJ. DoubletFinder: Doublet Detection in Single-Cell RNA Sequencing Data Using Artificial Nearest Neighbors. Cell Syst 2019; 8: 329–337 e324.

21. Guo M, Morley MP, Jiang C, Wu Y, Li G, Du Y, et al. Guided construction of single cell reference for human and mouse lung. Nat Commun 2023; 14: 4566.

22. Tabula Muris C. A single-cell transcriptomic atlas characterizes ageing tissues in the mouse. Nature 2020; 583: 590–595.

23. Strunz M, Simon LM, Ansari M, Kathiriya JJ, Angelidis I, Mayr CH, et al. Alveolar regeneration through a Krt8+ transitional stem cell state that persists in human lung fibrosis. Nat Commun 2020; 11: 3559.

24. Redente EF, Keith RC, Janssen W, Henson PM, Ortiz LA, Downey GP, et al. Tumor necrosis factor-alpha accelerates the resolution of established pulmonary fibrosis in mice by targeting profibrotic lung macrophages. Am J Respir Cell Mol Biol 2014; 50: 825–837.

25. Jin S, Plikus MV, Nie Q. CellChat for systematic analysis of cell-cell communication from single-cell transcriptomics. Nat Protoc 2025; 20: 180–219.

26. Liu H, Coarfa C, Charania AN, Larson-Casey JL, Rosas IO, He C. Secreted Phosphoprotein 1 in Lung Diseases. Metabolites 2025; 15: 365.

27. Laddha AP, Kulkarni YA. VEGF and FGF-2: Promising targets for the treatment of respiratory disorders. Respir Med 2019; 156: 33–46.

28. Antoniades HN, Bravo MA, Avila RE, Galanopoulos T, Neville-Golden J, Maxwell M, et al. Platelet-derived growth factor in idiopathic pulmonary fibrosis. J Clin Invest 1990; 86: 1055–1064.

29. Bonner JC. Regulation of PDGF and its receptors in fibrotic diseases. Cytokine Growth Factor Rev 2004; 15: 255–273.

30. Baxter SK, Irizarry-Caro RA, Vander Heiden JA, Arron JR. Breaking the cycle: should we target inflammation, fibrosis, or both? Front Immunol 2025; 16: 1569501.

31. Stawski L, Trojanowska M. Oncostatin M and its role in fibrosis. Connect Tissue Res 2019; 60: 40–49.

32. Kim SH, Nouws J, Ruwisch J, Woodard GA, Cooley J, Khoury J, et al. Dysregulated alveolar type 2 epithelial cell proteostasis promotes fibrogenic macrophage migration inhibitory factor-CD74 signaling. Sci Transl Med 2025; 17: eadr2277.

33. Kim BM, Lee YJ, Choi YH, Park EM, Kang JL. Gas6 Ameliorates Inflammatory Response and Apoptosis in Bleomycin-Induced Acute Lung Injury. Biomedicines 2021; 9.

34. Liu L, Wu P, Wei Y, Lu M, Ge H, Wang P, et al. TWEAK-Fn14 signaling protects mice from pulmonary fibrosis by inhibiting fibroblast activation and recruiting pro-regenerative macrophages. Cell Rep 2025; 44: 115220.

35. Dai Z, Song G, Balakrishnan A, Yang T, Yuan Q, Mobus S, et al. Growth differentiation factor 11 attenuates liver fibrosis via expansion of liver progenitor cells. Gut 2020; 69: 1104–1115.

36. Sawant H, Borthakur A. Disease-Specific Novel Role of Growth Differentiation Factor 15 in Organ Fibrosis. Int J Mol Sci 2025; 26.

37. Ye Q, Taleb SJ, Zhao J, Zhao Y. Emerging role of BMPs/BMPR2 signaling pathway in treatment for pulmonary fibrosis. Biomed Pharmacother 2024; 178: 117178.

38. Jin H, Mou J, Zhu H, Liu K, Zhang M, Zhang Z, et al. Lineage tracing reveals the origins and dynamics of macrophages in lung injury and repair. Cell Discov 2026; 12: 3.

39. Yadav P, Gomez Ortega J, Dabral P, Tamaki W, Chien C, Chang KC, et al. Myeloid-mesenchymal crosstalk drives ARG1-dependent profibrotic metabolism via ornithine in lung fibrosis. J Clin Invest 2025; 135.

40. Fabre T, Barron AMS, Christensen SM, Asano S, Bound K, Lech MP, et al. Identification of a broadly fibrogenic macrophage subset induced by type 3 inflammation. Sci Immunol 2023; 8: eadd8945.

41. King EM, Zhao Y, Moore CM, Steinhart B, Anderson KC, Vestal B, et al. Gpnmb and Spp1 mark a conserved macrophage injury response masking fibrosis-specific programming in the lung. JCI Insight 2024; 9.

42. Cui H, Banerjee S, Xie N, Hussain M, Jaiswal A, Liu H, et al. TREM2 promotes lung fibrosis via controlling alveolar macrophage survival and pro-fibrotic activity. Nature Communications 2025; 16.

43. Tsukui T, Wolters PJ, Sheppard D. Alveolar fibroblast lineage orchestrates lung inflammation and fibrosis. Nature 2024; 631: 627–634.

44. Crestani B, Marchand-Adam S, Quesnel C, Plantier L, Borensztajn K, Marchal J, et al. Hepatocyte growth factor and lung fibrosis. Proc Am Thorac Soc 2012; 9: 158–163.

45. Panganiban RA, Day RM. Hepatocyte growth factor in lung repair and pulmonary fibrosis. Acta Pharmacol Sin 2011; 32: 12–20.

46. Bocchi E, Pitozzi V, Pontis S, Caruso PL, Beghi S, Caputi M, et al. Investigation of Aberrant Basaloid Cells in a Rat Model of Lung Fibrosis. Front Biosci (Landmark Ed) 2024; 29: 305.

47. Chakraborty A, Mastalerz M, Ansari M, Schiller HB, Staab-Weijnitz CA. Emerging Roles of Airway Epithelial Cells in Idiopathic Pulmonary Fibrosis. Cells 2022; 11.

48. Ting C, Konopka K, Benedeck RE, Riches DWH, Redente EF, Oldham JM, et al. Biomarkers Unveil Insights into Pathology of Transitional Epithelial States in Pulmonary Fibrosis. Am J Respir Crit Care Med 2024; 210: 687–690.

49. Wang F, Ting C, Riemondy KA, Douglas M, Foster K, Patel N, et al. Regulation of epithelial transitional states in murine and human pulmonary fibrosis. J Clin Invest 2023; 133.

50. Dorry SJ, Ansbro BO, Ornitz DM, Mutlu GM, Guzy RD. FGFR2 Is Required for AEC2 Homeostasis and Survival after Bleomycin-induced Lung Injury. Am J Respir Cell Mol Biol 2020; 62: 608–621.

51. Shimbori C, El Agha E. Good Things Come in 2s: Type 2 Alveolar Epithelial Cells and Fibroblast Growth Factor Receptor 2. Am J Respir Cell Mol Biol 2020; 62: 543–545.

52. Francois A, Chatelus E, Wachsmann D, Sibilia J, Bahram S, Alsaleh G, et al. B lymphocytes and B-cell activating factor promote collagen and profibrotic markers expression by dermal fibroblasts in systemic sclerosis. Arthritis Res Ther 2013; 15: R168.

53. Horowitz JC, Thannickal VJ. Mechanisms for the Resolution of Organ Fibrosis. Physiology (Bethesda) 2019; 34: 43–55.

54. Tsukui T, Sun KH, Wetter JB, Wilson-Kanamori JR, Hazelwood LA, Henderson NC, et al. Collagen-producing lung cell atlas identifies multiple subsets with distinct localization and relevance to fibrosis. Nat Commun 2020; 11: 1920.

55. Planer JD, Morrisey EE. After the Storm: Regeneration, Repair, and Reestablishment of Homeostasis Between the Alveolar Epithelium and Innate Immune System Following Viral Lung Injury. Annu Rev Pathol 2023; 18: 337–359.

56. Song L, Li K, Chen H, Xie L. Cell Cross-Talk in Alveolar Microenvironment: From Lung Injury to Fibrosis. Am J Respir Cell Mol Biol 2024; 71: 30–42.

57. Wang Y, Wang L, Ma S, Cheng L, Yu G. Repair and regeneration of the alveolar epithelium in lung injury. FASEB J 2024; 38: e23612.

58. Redente EF, Black BP, Backos DS, Bahadur AN, Humphries SM, Lynch DA, et al. Persistent, Progressive Pulmonary Fibrosis and Epithelial Remodeling in Mice. Am J Respir Cell Mol Biol 2021; 64: 669–676.

59. Ban J, Qian J, Zhang C, Li J. Recent advances in TAM mechanisms in lung diseases. J Transl Med 2025; 23: 479.

60. Suzuki A, Sakamoto K, Nakahara Y, Enomoto A, Hino J, Ando A, et al. BMP3b Is a Novel Antifibrotic Molecule Regulated by Meflin in Lung Fibroblasts. Am J Respir Cell Mol Biol 2022; 67: 446–458.

61. Takenouchi Y, Kitakaze K, Tsuboi K, Okamoto Y. Growth differentiation factor 15 facilitates lung fibrosis by activating macrophages and fibroblasts. Exp Cell Res 2020; 391: 112010.

62. Cahill EF, Kennelly H, Carty F, Mahon BP, English K. Hepatocyte Growth Factor Is Required for Mesenchymal Stromal Cell Protection Against Bleomycin-Induced Pulmonary Fibrosis. Stem Cells Transl Med 2016; 5: 1307–1318.

63. Gazdhar A, Temuri A, Knudsen L, Gugger M, Schmid RA, Ochs M, et al. Targeted gene transfer of hepatocyte growth factor to alveolar type II epithelial cells reduces lung fibrosis in rats. Hum Gene Ther 2013; 24: 105–116.

64. Mizuno S, Matsumoto K, Li MY, Nakamura T. HGF reduces advancing lung fibrosis in mice: a potential role for MMP-dependent myofibroblast apoptosis. FASEB J 2005; 19: 580–582.

65. Shukla MN, Rose JL, Ray R, Lathrop KL, Ray A, Ray P. Hepatocyte growth factor inhibits epithelial to myofibroblast transition in lung cells via Smad7. Am J Respir Cell Mol Biol 2009; 40: 643–653.

66. Kinoshita K, Aono Y, Azuma M, Kishi J, Takezaki A, Kishi M, et al. Antifibrotic effects of focal adhesion kinase inhibitor in bleomycin-induced pulmonary fibrosis in mice. Am J Respir Cell Mol Biol 2013; 49: 536–543.

67. Lagares D, Busnadiego O, Garcia-Fernandez RA, Kapoor M, Liu S, Carter DE, et al. Inhibition of focal adhesion kinase prevents experimental lung fibrosis and myofibroblast formation. Arthritis Rheum 2012; 64: 1653–1664.

68. Brownfield DG, de Arce AD, Ghelfi E, Gillich A, Desai TJ, Krasnow MA. Alveolar cell fate selection and lifelong maintenance of AT2 cells by FGF signaling. Nat Commun 2022; 13: 7137.

69. Fortier SM, Penke LR, King D, Pham TX, Ligresti G, Peters-Golden M. Myofibroblast dedifferentiation proceeds via distinct transcriptomic and phenotypic transitions. JCI Insight 2021; 6.

70. Guzy RD, Li L, Smith C, Dorry SJ, Koo HY, Chen L, et al. Pulmonary fibrosis requires cell-autonomous mesenchymal fibroblast growth factor (FGF) signaling. J Biol Chem 2017; 292: 10364–10378.

71. Wiley SR, Cassiano L, Lofton T, Davis-Smith T, Winkles JA, Lindner V, et al. A novel TNF receptor family member binds TWEAK and is implicated in angiogenesis. Immunity 2001; 15: 837–846.

72. Zhang Y, Zeng W, Xia Y. TWEAK/Fn14 axis is an important player in fibrosis. J Cell Physiol 2021; 236: 3304–3316.

73. Gomez IG, Roach AM, Nakagawa N, Amatucci A, Johnson BG, Dunn K, et al. TWEAK-Fn14 Signaling Activates Myofibroblasts to Drive Progression of Fibrotic Kidney Disease. J Am Soc Nephrol 2016; 27: 3639–3652.

74. Burkly LC. Regulation of Tissue Responses: The TWEAK/Fn14 Pathway and Other TNF/TNFR Superfamily Members That Activate Non-Canonical NFkappaB Signaling. Front Immunol 2015; 6: 92.

75. Peters-Golden M, Fortier SM. Mechanistic basis for the antifibrotic actions of cAMP-based therapies. Eur Respir Rev 2026; 35.

76. Huang SK, White ES, Wettlaufer SH, Grifka H, Hogaboam CM, Thannickal VJ, et al. Prostaglandin E(2) induces fibroblast apoptosis by modulating multiple survival pathways. FASEB J 2009; 23: 4317–4326.

77. Reininger D, Wolf F, Mayr CH, Wespel SL, Laufhaeger N, Geillinger-Kastle K, et al. Insights into the Cellular and Molecular Mechanisms behind the Antifibrotic Effects of Nerandomilast. Am J Respir Cell Mol Biol 2025; 73: 700–712.

78. Nathan SD, Smith P, Deng C, De Salvo M, Wuyts W, Pavie-Gallegos J, et al. Inhaled Treprostinil for Idiopathic Pulmonary Fibrosis. N Engl J Med 2026.

